# From anti-fungal to potential neurotherapeutic: Posaconazole as an effective inhibitor of cellular TDP-43 pathology

**DOI:** 10.64898/2026.06.01.728552

**Authors:** Noah Nathan Kochen, Sophia Zafari, Aimee Renaud, Nathan Schneider, Nagamani Vunnam, Elly E. Liao, James R. Dutton, Anthony R. Braun, Jonathan N. Sachs

**Affiliations:** Department of Biomedical Engineering, University of Minnesota, Minneapolis, MN 55455; Stem Cell Institute, University of Minnesota Medical School, Minneapolis, MN 55455

## Abstract

Recently, we showed that ketoconazole, a known anti-fungal inhibitor of CYP51, stabilized TAR DNA-binding protein 43 (TDP-43) native self-interactions, reduced TDP-43 pathology and rescued TDP-43-induced SREBP2 downregulation. Despite its promising effects, ketoconazole is not viable for repurposing for ALS due to liver toxicity side effects that occur when orally delivered. To address this, we tested the activities of seven additional known azole-based CYP51 inhibitors in order identify a viable alternative to ketoconazole. Using our established TDP-43 mislocalization and aggregation assay in HEK293T cells, we identified posaconazole, an FDA-approved, CNS-penetrant and orally delivered anti-fungal, as the strongest inhibitor of TDP-43 pathology. Posaconazole was able to reduce insoluble TDP-43 and restore SREBP2 levels, outperforming ketoconazole. Mechanism of action (MOA) experiments suggest posaconazole is able to outperform ketoconazole by inducing a significantly stronger activation of autophagy and upregulation of heat shock proteins known to clear TDP-43. Further MOA experiments show that the effects of posaconazole on TDP-43 are dependent on its known ability to lower cellular cholesterol levels. By correlating our experimental results on the eight CYP51 inhibitors tested, we show that predicted affinity towards human CYP51 strongly correlates with the inhibitors’ ability to lower TDP-43 aggregation and mislocalization. Finally, we tested posaconazole in a low dose sodium arsenite ALS model in iPSC-derived motor neurons, showing that it is efficacious at inhibiting TDP-43 pathology in the nanomolar range. Altogether, these results support the repurposing of posaconazole for ALS/FTD as a means to prevent TDP-43 pathology.

## Introduction

TAR DNA-binding protein 43 (TDP-43) is a crucial protein involved in the regulation of hundreds of RNA targets, cellular stress responses and axonal transport of mRNAs (*1*). Its involvement in amyotrophic lateral sclerosis (ALS), frontotemporal dementia (FTD) and Alzheimer’s disease pathology has been attributed to a dual loss of native nuclear function and a gain of toxic function triggered by its mislocalization into the cytoplasm, where it can aggregate into insoluble amyloid fibrils (*2, 3*). There are currently no approved treatments that directly address TDP-43 pathology, despite it being present in roughly 97% of ALS cases and 45% of FTD cases (*4*).

We recently developed a high-throughput screening platform capable of measuring functional TDP-43 nuclear species and their self-interactions in living cells, which we used to screen the FDA-approved Selleck library (*5*). We identified the hit compound ketoconazole, a known azole-containing inhibitor of both fungal and human 14α-lanosterol demethylase (CYP51), a P450 cytochrome enzyme that demethylates lanosterol in the first step of its conversion into cholesterol (in humans) or ergosterol (in fungi) (*6, 7*). Ketoconazole increased native nuclear TDP-43 interactions, rescued TDP-43 mislocalization and aggregation, and improved the health of neurons overexpressing TDP-43 (*5*). In addition, we showed that ketoconazole’s stabilization of TDP-43 self-interactions correlated with an increase in mRNA levels of SREBP2, the master regulator of the cholesterol biosynthesis pathway. Interestingly, ketoconazole’s effects were blocked when co-treating with the SCAP inhibitor betulin, which sequesters SREBP2 in the endoplasmic reticulum and blocks its downstream pathway (*5*). These preliminary mechanistic experiments pointed to the modulation of CYP51, SREBP2 and the cholesterol pathway (which is known to be disrupted in ALS and TDP-43 pathology) as the most likely mechanism of action linking ketoconazole and TDP-43.

Indeed, there is evidence that TDP-43 is actively involved in regulating the cholesterol pathway by binding to mRNAs of SREBP2 and its downstream regulated enzymes involved in sterol synthesis, such as CYP51 (*8, 9*). Interestingly, disruption of TDP-43 function via overexpression or knockout leads to major disruptions in the homeostatic expression patterns of genes in this pathway, with SREBP2 being one of the most affected genes via downregulation (*8, 9*). Maintenance of SREBP2 levels is crucial for cholesterol homeostasis in the CNS, as it regulates myelin production, synapse integrity, proteostasis and stress responses (*10-13*). Though not explored yet, the known involvement of cholesterol levels in modulating the heat shock protein response and autophagy (*14, 15*)—two major pathways involved in preventing and clearing TDP-43 pathological aggregation—suggests an intricate relationship between cholesterol homeostasis and TDP-43 proteostasis that may have important implications for developing new therapeutics for ALS/FTD. In our previous work, we showed that ketoconazole was able to rescue TDP-43-induced downregulation of SREBP2, but this effect was only partial (*5*).

A patent filed in 2019 by Yumanity Therapeutics proposed inhibition of CYP51 as a therapeutic target for TDP-43-associated diseases such as ALS and FTD (*16*). The authors identified the anti-fungal fluconazole as an inhibitor of toxicity in human WT TDP-43 overexpressing yeast. In addition, they report that two previously published non-azole based CYP51 inhibitors can ameliorate the survival of primary rat cortical neurons transfected with WT or mutant Q331K TDP-43 (*16*). Nevertheless, this patent lacked 1) evidence that azole-based anti-fungals have an effect in human cells, 2) a direct comparison between potency towards CYP51 and efficacy in reducing TDP-43 pathology; and 3) any mechanistic explanation for the rescue of TDP-43 pathology by CYP51 inhibition (*16*). Since the submission of this patent, Yumanity Therapeutics reverse merged with Kineta and sold its therapeutic candidates to Johnson & Johnson (*17*). To the best of our knowledge, no research articles, besides our previous (*5*) and current work, have been publicly published by Yumanity Therapeutics or other groups linking CYP51 inhibition to TDP-43 before or since the deposition of this patent.

Importantly, ketoconazole’s use in the United States has been severely restricted to topical treatments as of 2013, as oral delivery can cause fatal liver toxicity (*18*). If ketoconazole acts by modulating the cholesterol pathway through inhibition of CYP51, we hypothesized that this therapeutic effect could be achieved with a different molecule in the family of anti-fungal azoles that has a better safety profile. Therefore, we set out to 1) validate anti-fungal azoles as a class of compounds with activity towards TDP-43; 2) identify a safe alternative molecule than our previously identified hit ketoconazole; and 3) elucidate whether the activity of anti-fungal azoles on TDP-43 is dependent on their inhibition of cholesterol synthesis. To answer these questions, we evaluated the efficacy of seven additional azoles with known activity towards CYP51 other than ketoconazole, namely: fenticonazole; itraconazole; clotrimazole; cyproconazole; fluconazole; voriconazole; and posaconazole. Using a combination of live-cell fluorescence imaging and biochemical characterization, we show that posaconazole, a recently developed triazole-based inhibitor, outperforms ketoconazole in reducing TDP-43 pathology, correlating with its enhanced ability to restore SREBP2 expression, upregulate heat shock proteins and activate autophagy. Using a cholesterol supplementation model, we demonstrate that the therapeutic effects of ketoconazole and posaconazole on TDP-43 are blocked when cellular cholesterol levels are increased. Using molecular docking and structural predictions, we show that activities of the azoles towards human CYP51 correlate with their effects on TDP-43 mislocalization and aggregation. Finally, we show that posaconazole is able to lower TDP-43-induced pathology in an iPSC-derived motor neuron model of ALS that displays TDP-43 pathology in response to low dose sodium arsenite. Altogether, our results position posaconazole as a candidate for drug repurposing for TDP-43-related diseases such as ALS and FTD.

## Results

### Known anti-fungal azole inhibitors of CYP51 rescue TDP-43 mislocalization and aggregation in HEK293T cells

Previously, we showed that ketoconazole, rescued sorbitol-induced TDP-43 mislocalization and aggregation in a dose dependent manner, suggesting a connection between CYP51 inhibition and rescue of TDP-43 pathology (*5*). In order to identify a safer and orally delivered alternative to ketoconazole, we evaluated a panel of 7 additional known anti-fungal azole inhibitors of CYP51 with structural similarity to ketoconazole (Fig. S1). First, we compared the activity of the compounds in our panel towards CYP51 using published fungal CYP51 inhibition studies in which DMSO, ketoconazole, fluconazole, posaconazole, itraconazole and voriconazole were tested (5 of the 8 inhibitors in our panel) (*6, 19, 20*). The fungal CYP51 enzymes tested in these studies (T. cruzi, A. fumigatus, C. albicans, S. parasitica) are similar to human CYP51 both structurally and in sequence, with an average RMSD relative to human CYP51 (PDB ID: 3LD6) of 1.80 ± 0.23 Å, ∼37% global sequence identity (exact amino acid matching), and ∼57% sequence identity (exact matching) within the CYP51 family’s conserved structural motifs (Fig. S2) (*21*). Fig. 1A shows the wide range in effects of these compounds towards fungal CYP51s. From these studies, posaconazole, a safe, FDA-approved and orally delivered anti-fungal derived from ketoconazole, was identified as the azole with highest activity, with an average residual CYP51 activity of 4.39%, ∼5x times lower than the average for ketoconazole across the four species (21.9%). In addition, itraconazole, another ketoconazole-derived compound showed stronger inhibition relative to ketoconazole (∼2x stronger inhibition). Clotrimazole displayed similar inhibition of fungal CYP51s as ketoconazole.

**Fig. 1.**
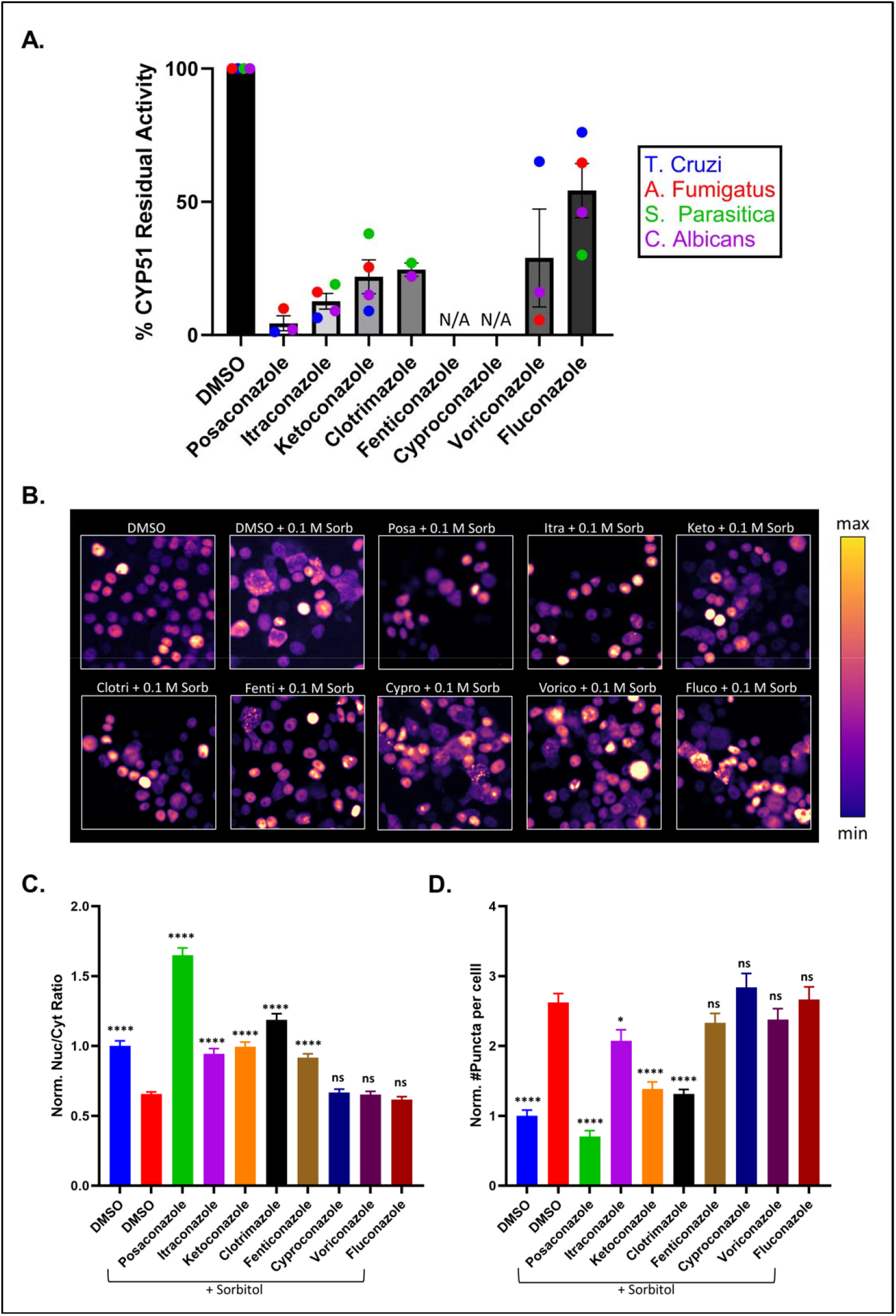
Sorbitol-induced TDP-43 mislocalization and aggregation is rescued by treatment with known anti-fungal CYP51 inhibitors. **(A)** %Residual activity of fungal CYP51s in the presence of azole inhibitors. Adapted from (*6, 19, 20*). **(B)** Live-cell images of TDP-43-mNg-expressing HEK293Ts treated with sorbitol and 10 μM azoles for 24 hours. Green fluorescence was mapped to a pseudo-color LUT. **(C-D)** Quantification of Nuc/Cyt ratios and puncta (data normalized to DMSO-only treated cells, blue bar). Data shown are mean ± SEM of N=3 independent experiments, analyzed via a one-way ANOVA with Bonferroni correction for multiple comparisons relative to DMSO/sorbitol-treated cells (red bar, *p < 0.05, ****p < 0.0001).

Next, we sought to compare the activity of these azoles towards reducing TDP-43 pathology. We used our established low hyperosmotic stress model, consisting of TDP-43-mNeonGreen expressing HEK293T cells treated with 0.1 M sorbitol and co-treated with the 8 azole inhibitors at 10 μM for 24 hours. As shown in Fig. S3, sorbitol treatment causes both TDP-43 cytoplasmic mislocalization and aggregation. Fig. 1B-D shows fluorescent images and quantification of the sorbitol-induced increase in mislocalization (measured via nuclear-to-cytoplasmic ratios, Nuc/Cyt) and cytoplasmic aggregation (measured via # of cytosolic puncta per cell) of TDP-43, which amounted to 35% and 162% that of untreated cells, respectively. Out of the 8 inhibitors, ketoconazole, posaconazole, fenticonazole, clotrimazole and itraconazole were able to significantly rescue mislocalization and/or aggregation relative to sorbitol/DMSO-only treated cells. Cyproconazole, fluconazole and voriconazole showed no statistically significant effects relative to sorbitol/DMSO-only treated cells. Out of the 7 additional azoles tested, itraconazole and clotrimazole, two safe and orally delivered anti-fungals, had a therapeutic effect comparable to ketoconazole. Interestingly, posaconazole was the only azole able to outperform ketoconazole, in agreement with its significantly stronger inhibition of CYP51 shown in Fig. 1A. In contrast, fluconazole, which had the lowest activity towards CYP51 in Fig. 1A, had no effect on TDP-43 relative to DMSO.

To further validate our findings on posaconazole, we examined sequential soluble/insoluble protein extractions under sorbitol and drug treatments (at 10 μM) using cells expressing TDP-43-mNg. Fig. S4A-B present immunoblots for TDP-43 (quantified in Fig. S4C-D) showing that the addition of sorbitol resulted in a 66% increase in insoluble TDP-43-mNg species relative to DMSO-only treated cells, agreeing with previous studies (*22, 23*). Both ketoconazole and posaconazole significantly decreased the insoluble TDP-43 fraction induced by sorbitol treatment, with posaconazole having a stronger effect (full recovery to no-sorbitol insoluble TDP-43-mNg levels). Figs. S4A and S4C show the quantification of soluble TDP-43 levels, which were slightly elevated in all conditions containing sorbitol, with no significant changes induced by the drugs relative to DMSO. Fig. S5 shows all western blots used for the characterization of insoluble and soluble TDP-43.

### Posaconazole fully rescues SREBP2 downregulation under TDP-43 overexpression

Having demonstrated that anti-fungal azoles are active towards TDP-43, and that posaconazole is more effective than ketoconazole in reducing TDP-43 pathology, we wanted to explore potential mechanisms of action that explain this increased efficacy. Given the known involvement of these azoles as CYP51 inhibitors, and the role of SREBP2 in TDP-43-related ALS, we examined the cholesterol biosynthesis pathway (*8, 9, 24*). TDP-43 pathology has been implicated in cholesterol biosynthesis deficits via the downregulation of the pathway’s master regulator SREBP2 (*8, 9*), which we previously showed can be partially rescued by ketoconazole (*5*). TDP-43 is a known regulator of cholesterol biosynthesis genes at the mRNA level, due to the widespread presence of GU-rich motifs with affinity towards TDP-43’s RNA-recognition motifs (RRMs). In addition, CYP51 inhibition is known to modulate the cholesterol pathway by inhibiting lanosterol demethylation in its conversion towards cholesterol (*7*). This has been shown to lead to an accumulation of lanosterol and reduction in cholesterol synthesis, activating an autoregulatory feedback loop that ultimately results in the upregulation of SREBP2 aimed at restoring sterol homeostasis (*25-27*). In our previous work, we proposed this mechanism for the observed effects of ketoconazole on SREBP2 expression (*5*).

First, we wanted to validate the downregulation of SREBP2 as a pathological phenotype in patients within the ALS/FTD spectrum, which are known to share TDP-43 pathology (*4*). We used the most recent NYGC ALS Consortium transcriptomics dataset, comprised of 695 donors in the ALS/FTD spectrum across 8 different CNS tissue types (*28, 29*). We first focused on the spinal cord, as it contains lower motor neurons that are heavily affected in ALS and FTD (*30*). As shown in Fig. 2A, SREBP2 expression was significantly reduced in cervical and lumbar spinal cord tissue, with the phenotype being stronger in the cervical spinal cord. Thoracic spinal cord samples showed a similar trend in a reduction in SREBP2 expression, but with no statistical significance. Interestingly, both the cervical and lumbar regions of the spinal cord contain anatomical enlargements which contain high numbers of lower motor neurons relative to the thoracic spinal cord (*31*). In addition to the spinal cord, Fig. S6 shows that SREBP2 is also significantly downregulated in the frontal cortex, which is affected in ALS/FTD, where its atrophy correlates with behavioral and cognitive decline (*32*). Other regions such as the temporal cortex, occipital cortex, medial motor cortex, lateral motor cortex, hippocampus and cerebellum had no significant changes in SREBP2 expression in ALS/FTD patients relative to healthy controls (Fig. S6), suggesting this phenotype may be restricted to the spinal cord.

**Fig. 2.**
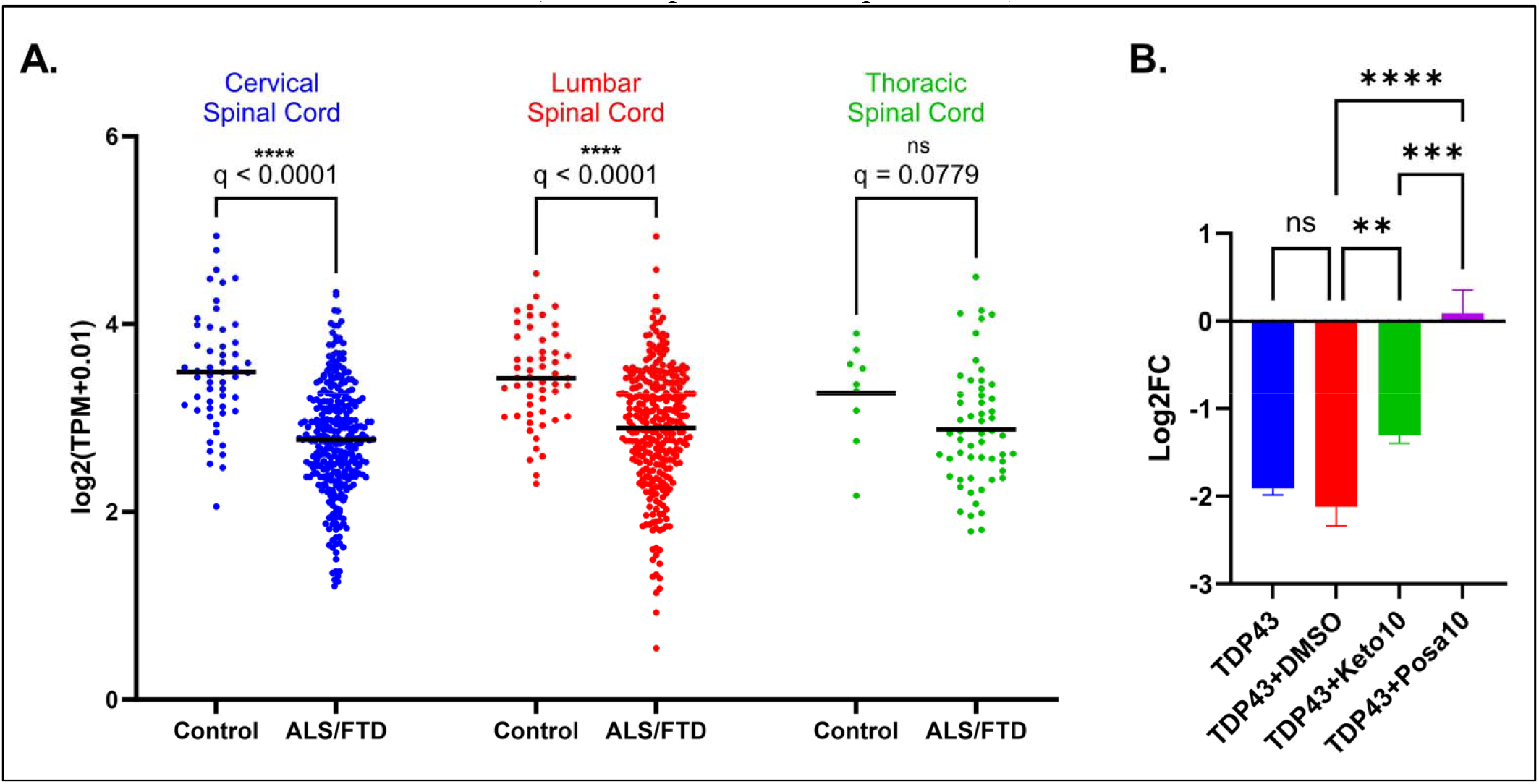
TDP-43-overexpression model recapitulates spinal cord SREBP2 downregulation phenotype in ALS/FTD patients and is rescued by posaconazole. **(A)** Transcripts per million (TPM) counts for SREBP2 mRNA in cervical, lumbar and thoracic spinal cord tissue of healthy controls and ALS/FTD patients extracted from the NYGC ALS Consortium dataset. See Fig. S6 for SREBP2 expression across all tissues characterized in this dataset. Data was analyzed via a two-way ANOVA with Benjamini-Hochberg False Discovery Rate (FDR) correction (****q < 0.0001, q-values equate to FDR-adjusted p-values). **(B)** RT-qPCR Log2 fold changes (Log2FC) for SREBP2 in TDP-43 overexpressing cells treated with DMSO and 10 μM drugs for 24 hours. Fold changes were calculated relative to untreated cells using GAPDH as a housekeeping control. Data shown in are mean ± SEM of N=3 independent samples, analyzed via a one-way ANOVA with Bonferroni correction for multiple comparisons (*p < 0.05, **p<0.01, ***p<0.001, ****p < 0.0001).

Given its higher predicted activity towards CYP51, we hypothesized that posaconazole would ameliorate the SREBP2 downregulation induced by TDP-43 overexpression to a greater extent than ketoconazole. Fig. 2B shows that TDP-43 overexpression led to a ∼2-fold reduction in SREBP2 mRNA, recapitulating results published by Ho et.al. and Egawa et.al. (*8, 9*). SREBP2 levels were not affected by DMSO treatment relative to untreated TDP-43 overexpressing cells but partially rescued by 10 μM ketoconazole (∼1-fold increase) and fully rescued by 10 μM posaconazole (∼2-fold increase) back to control cells’ levels. These results correlate with cellular cholesterol measurements in cells treated with 10 μM ketoconazole and posaconazole for 12 hours, where we observed a more significant decrease with posaconazole (∼30% vs. 20% reduction relative to DMSO-only, Fig. S7A). At 24 hours, both drugs lowered cholesterol levels by ∼40% of DMSO-only treated cells, consistent with previous reports of their ability to inhibit cholesterol synthesis (Fig. S7B) (*33*). The fact that posaconazole is able to decrease cholesterol levels further than ketoconazole at 12 hours, but shows no difference at 24 hours, suggests that posaconazole can modulate this pathway more rapidly and that negative feedback regulation of the cholesterol pathway is likely at play.

### Posaconazole outperforms ketoconazole in upregulating heat shock proteins and activating autophagy

Although we observed a correlation between modulation of the cholesterol pathway and reduction in TDP-43 pathology, this does not provide a clear molecular mechanism explaining posaconazole’s stronger effect in reducing TDP-43 pathology relative to ketoconazole. After thoroughly searching the literature, we rationalized that inducing cholesterol depletion, which is sensed by the cells as nutrient starvation (*34*), can lead to the activation of heat shock proteins and autophagy—two pathways known to clear insoluble TDP-43 (*14, 15*). In addition, lanosterol accumulation, which itself modulates proteostasis via direct dissolution of protein aggregates, can lead to the upregulation of heat shock proteins (*35, 36*) which are reduced by TDP-43 mutations and overexpression (*37, 38*). Thus, we hypothesized that given its more potent modulation of the cholesterol pathway, posaconazole would lead to a stronger activation of TDP-43-related heat shock proteins and autophagy known to reduce TDP-43 pathological burden.

Fig. 3A presents RT-qPCR data showing that TDP-43 overexpression results in a global downregulation of heat shock proteins with known TDP-43 involvement (HSPB5, HSPB1, HSPB8, DNAJB2, HSP90A, HSP90B and HSP70); with no significant further changes induced by DMSO treatment. In contrast, 10 μM ketoconazole and posaconazole were able to significantly upregulate the levels of HSPB5, HSPB8, DNAJB2 and HSPB1, with posaconazole having a greater overall effect relative to ketoconazole. In addition, only posaconazole increased the levels of HSP90A and HSP90B, and, intriguingly, reduced the levels of HSP70 relative to DMSO-treated TDP-43 overexpressing cells, which is addressed later in the Discussion.

**Fig. 3.**
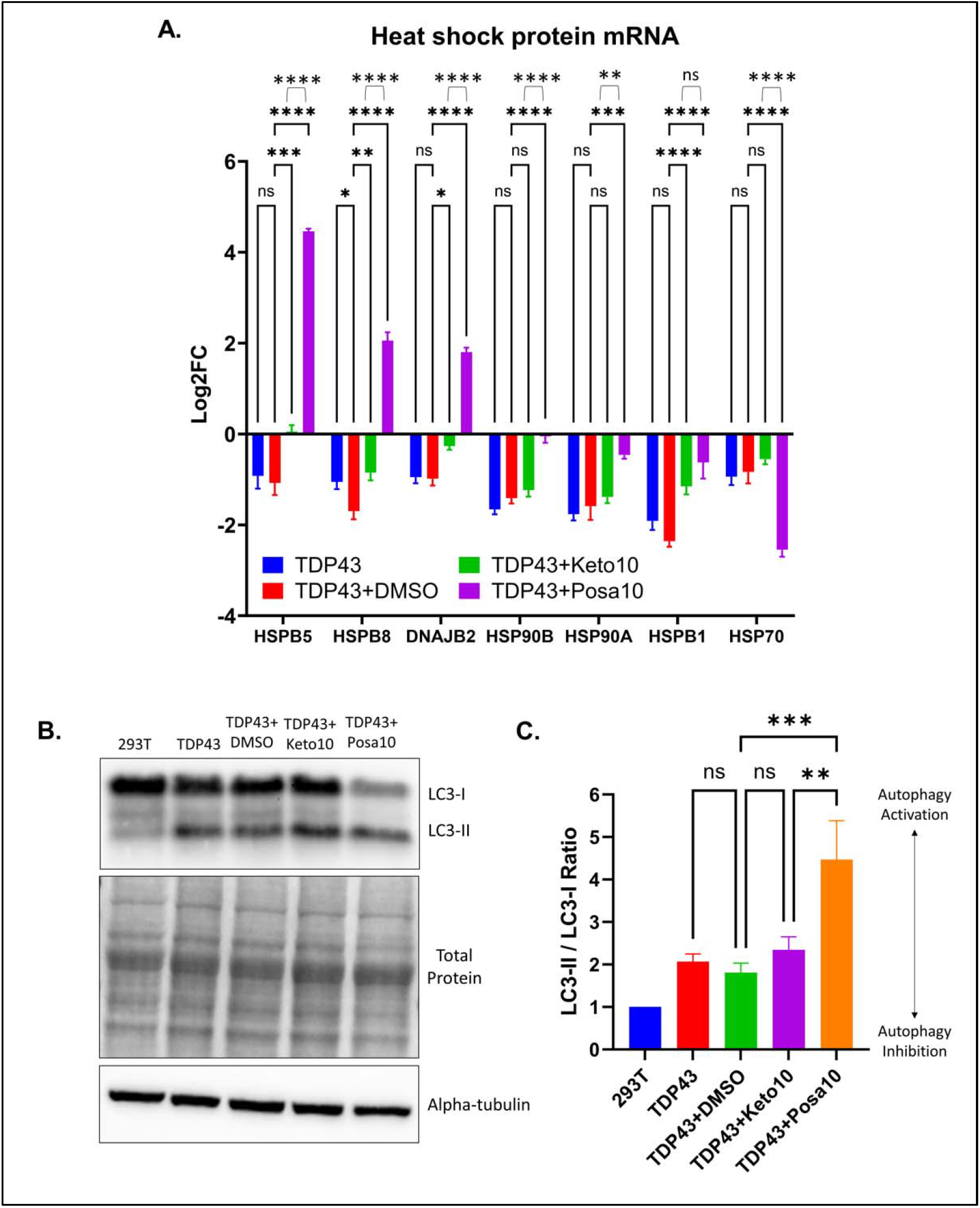
Posaconazole strongly increases levels of heat shock proteins and autophagy in TDP-43-overexpressing cells. **(A)** RT-qPCR Log2 fold changes (Log2FC) for panel of heat shock proteins in TDP-43 overexpressing cells treated with DMSO and 10 μM drugs for 24 hours. Fold changes were calculated relative to untreated cells using GAPDH as a housekeeping control. Data shown are mean ± SEM of N=3 independent samples, analyzed via a two-way ANOVA with Bonferroni correction for multiple comparisons (*p < 0.05, **p<0.01, ***p<0.001, ****p < 0.0001). **(B)** Representative western blot for LC3 in control cells and TDP-43 overexpressing cells treated with DMSO, 10 μM ketoconazole and 10 μM posaconazole for 24 hours (see Fig. S8 for N=2-9 blots) normalized to untreated cells (single untreated control per independent experiment). Alpha-tubulin was used as a loading control. **(C)** Quantification of LC3-II/LC3-I ratios. Data shown are mean ± SEM of N=9 independent samples, analyzed via a one-way ANOVA with Bonferroni correction for multiple comparisons (*p < 0.05, **p<0.01, ***p<0.001, ****p < 0.0001).

For autophagy, we probed for the ratio of LC3-II/LC3-I under TDP-43 overexpression conditions with DMSO, ketoconazole and posaconazole treatments. The levels of LC3-I (cytosolic form) and LC3-II (lipidated form that associates with autophagosome membranes) serve as an indicator of autophagy, where a relative increase in the LC3-II/LC3-I ratio indicates activation of this pathway (*39*). Fig. 3B-C shows that TDP-43 overexpression led to a mild increase in the basal ratio of LC3-II/LC3-I in untransfected cells, which is commonly observed under proteostatic stress conditions induced by aggregation-prone proteins (*40*). Only posaconazole induced a statistically significant increase in the ratio after 24 hours of treatment relative to DMSO-treated cells overexpressing TDP-43 (*39*). Ketoconazole mildly increased the ratio, but with no statistical significance relative to DMSO. Fig. S8 shows all western blots and the levels of LC3-I and LC3-II normalized to untransfected cells, showing that the increase in ratio observed for posaconazole was mainly driven by a reduction in LC3-I levels. Together with posaconazole’s ability to reduce insoluble TDP-43, the significant increase in LC3-II/LC3-I and reduction in LC3-I suggests an efficient conversion of LC3-I into LC3-II and subsequent autophagic clearance. Altogether, these results suggest posaconazole’s greater effect on TDP-43 pathology relative to ketoconazole are in part explained by a stronger activation of heat shock proteins and autophagy triggered by its strong modulation of the cholesterol pathway.

### Cholesterol supplementation blocks posaconazole’s and ketoconazole’s effects on TDP-43

To further demonstrate that ketoconazole and posaconazole’s mechanism of action is mediated by modulation of the cholesterol pathway, we incorporated cholesterol supplementation in our TDP-43 mislocalization and aggregation assay. The reason we performed these supplementation experiments was to further strengthen the likelihood that cholesterol depletion is at play in mediating the effects of ketoconazole and posaconazole on TDP-43. First, as shown in Fig. S9, we validated that supplementation with preloaded 6:1 methyl-beta cyclodextrin (MBCD):cholesterol complexes resulted in an increase in total cholesterol levels. Fig. 4A-C shows that cholesterol supplementation does not affect TDP-43 basal or sorbitol-induced mislocalization and aggregation, but that it significantly blocks the rescuing effect of ketoconazole and posaconazole, reversing their therapeutic effects near sorbitol/DMSO-only treated cells.

**Fig. 4.**
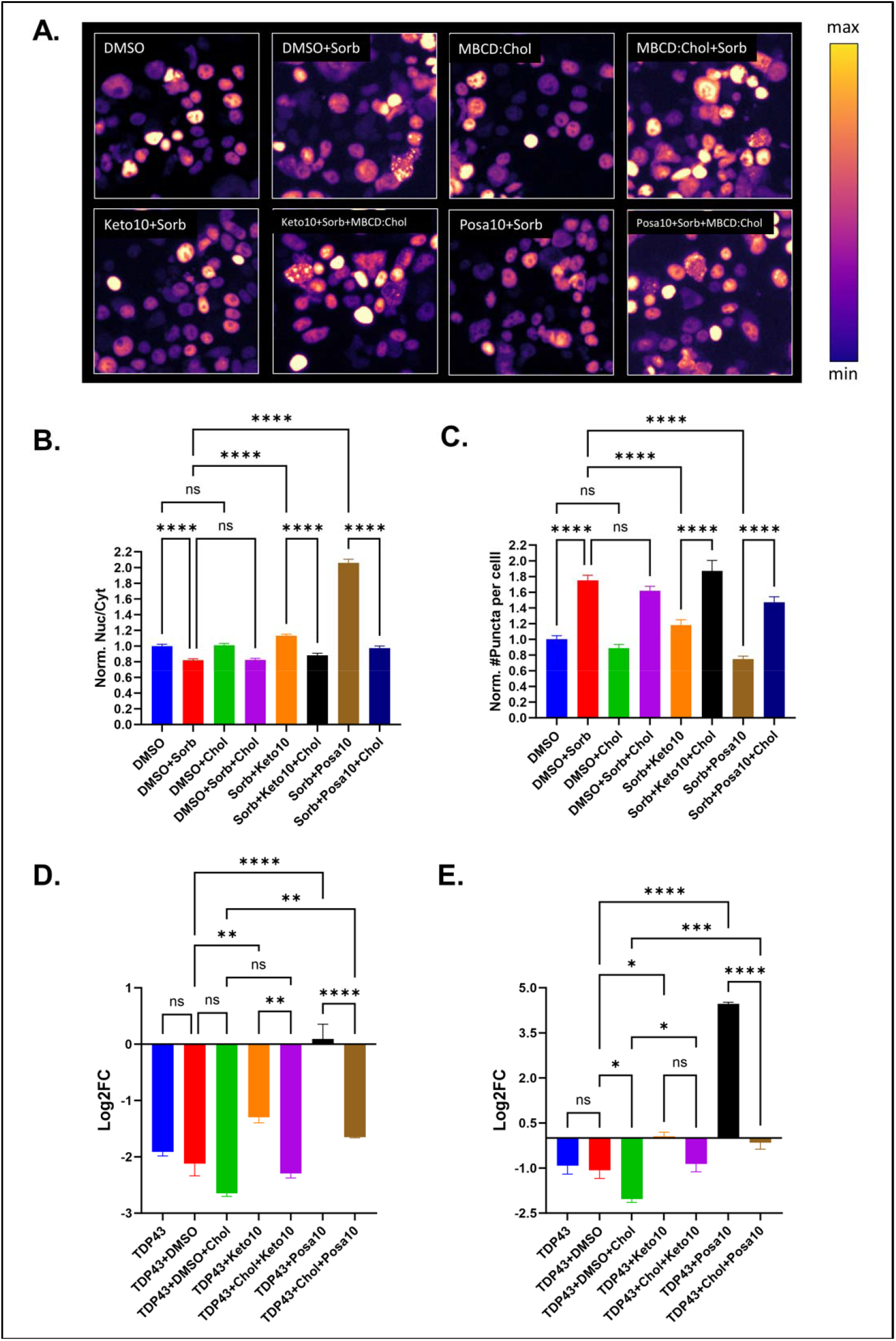
Cholesterol supplementation blocks effects of ketoconazole and posaconazole on TDP-43. **(A)** Live-cell images of TDP-43-mNg-expressing HEK293Ts treated with sorbitol and 10 μM azoles for 24 hours -/+ 100 μM cholesterol. **(B-C)** Quantification of Nuc/Cyt ratios and puncta per cell (data normalized to DMSO-only treated cells, blue bar). **(D-E)** Log2FC for **(D)** SREBP2 and **(E)** HSPB5 in TDP-43 overexpressing cells treated with 10 μM azoles for 24 hours -/+ 100 μM cholesterol. Fold changes were calculated relative to untreated cells using GAPDH as a housekeeping control. Data shown are mean ± SEM of N=3 independent experiments, analyzed via a one-way ANOVA with Bonferroni correction for multiple comparisons (*p < 0.05, **p<0.01, ***p<0.001, ****p < 0.0001).

As a final validation that the cholesterol pathway is at play in mediating the effects of ketoconazole and posaconazole, we tested whether cholesterol supplementation would block the observed upregulation of SREBP2 and the most upregulated heat shock protein (HSPB5) in our RT-qPCR assay. Fig. 4D-E, shows that cholesterol supplementation alone led to a decrease in both SREBP2 levels (consistent with the cholesterol pathway autoregulatory feedback loop) and HSPB5 levels relative to DMSO-only treated cells. In addition, Fig. 4D-E shows that the upregulation of SREBP2 and HSPB5 induced by the two azoles is significantly blocked by cholesterol supplementation. Altogether, these data strongly suggest that the effects of ketoconazole and posaconazole towards TDP-43 are mediated by the modulation of cellular cholesterol levels by controlling the levels of SREBP2 and HSPB5.

### Predicted azole inhibitor affinity towards human CYP51 correlates with TDP-43 cellular effects

An important question that we did not address in our cellular experiments is whether affinity towards human CYP51 would predict the effects of the anti-fungal azoles on TDP-43. This is crucial, as such an observation would more directly implicate the inhibition of this enzyme with a reduction in TDP-43 pathology. The binding affinities and inhibitory activities of the 8 azole inhibitors have been studied in the literature; however, most studies have focused on a limited number of these compounds or explored their activities towards fungal CYP51’s instead of the human ortholog (as shown in Fig. 1A). Thus, we turned to molecular docking and AI-based structural prediction tools (Boltz-2) to predict the activity of our azole panel towards human CYP51. First, we leveraged the structure of human CYP51 co-crystallized with ketoconazole (PDB ID: 3LD6) (*7*) to perform *in silico* molecular docking of the 8 inhibitors, ensuring their known heme-azole binding mode via a metal coordination constraint (imidazole or triazole group interacting with the heme iron). We confirmed that the metal coordination constraint resulted in a similar ketoconazole pose as the co-crystallized bound ligand (Fig. 5A; RMSD = 0.68 Å). Next, we docked the 7 other inhibitors (and DMSO as a negative control) to human CYP51 (Glide docking scores and ligand-protein interaction diagrams shown in Fig. S10). Out of the 8 azoles, posaconazole resulted in the best docking score (-10.518 vs. -8.350 for ketoconazole), and fluconazole had the worst score (-4.073 relative vs. -8.350 for ketoconazole).

**Fig. 5.**
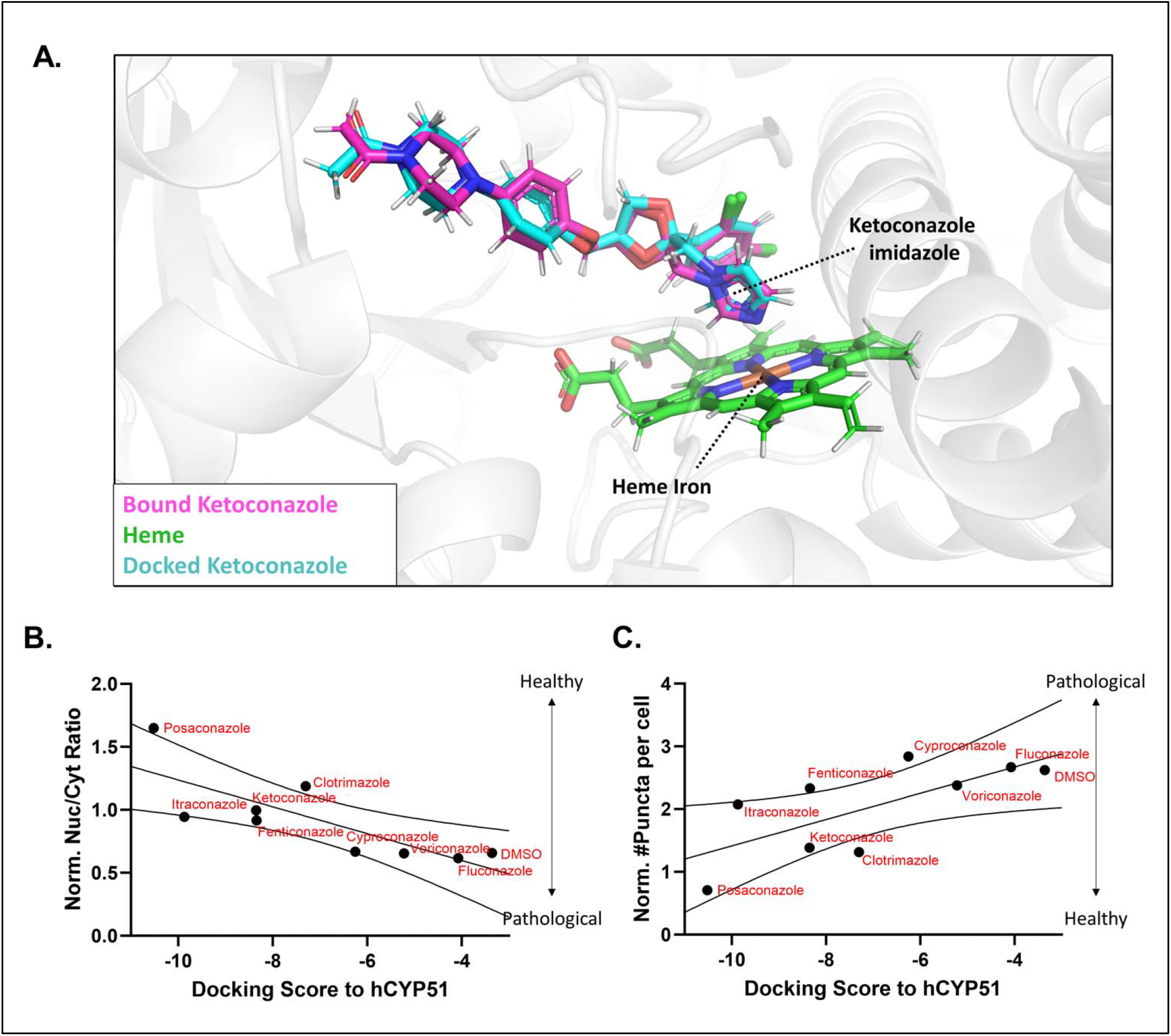
Molecular docking to human CYP51 predicts anti-fungal azole effects on TDP-43 pathology. **(A)** Comparison of human CYP51-bound ketoconazole (magenta) and docked ketoconazole (cyan) relative to co-crystallized heme group (PDB ID: 3LD6). RMSD between docked and co-crystallized ketoconazole was 0.68 Å). **(B)** Correlation between docking score and Nuc/Cyt ratios for DMSO and azole inhibitors (** p = 0.0111). **(C)** Correlation between docking score and # of puncta for DMSO and azole inhibitors (*p = 0.0308).

As an orthogonal validation of our docking studies, we used the recently developed AI-based Boltz-2 model for protein-ligand structural prediction to predict affinities for each inhibitor to human CYP51 without implementing any protein-ligand binding constraints, as only protein sequence and the SMILES representation of the 8 inhibitors are used as inputs (*41*). As shown in Fig. S11A, Boltz-2 was able to predict the azole-iron coordination binding mode for all inhibitors (RMSD = 0.531 Å for CYP51-ketoconazole complex relative to PDB ID: 3LD6). Out of the 8 azoles, posaconazole had the best predicted affinity in log_10_(IC50 μM) units (-1.71 vs. -1.26 for ketoconazole, Fig. S11B), and fluconazole had the worst (0.519 vs. -1.26 for ketoconazole, Fig. S11B), in agreement with the Glide docking results (linear correlation between docking and Boltz-2 predictions shown in Fig. S11C).

Next, we correlated human CYP51 docking scores to TDP-43 mislocalization and aggregation under sorbitol treatment for each of the tested inhibitors and DMSO. Fig. 5B-C shows that strong docking scores (more negative) significantly correlated with higher Nuc/Cyt ratios (lower mislocalization) and lower puncta per cell (lower aggregation). Notably, posaconazole, which had the strongest effects on mislocalization and aggregation, also had the best scores (both Glide and Boltz-2 predicted affinity). Conversely, fluconazole, a known weak inhibitor of CYP51, had the worst docking score and predicted affinity of all inhibitors and did not rescue mislocalization or aggregation at the tested concentration. Altogether, these correlations support our hypothesis that increased anti-fungal azole activity towards human CYP51 is sufficient to explain their therapeutic effects on TDP-43 in HEK293T cells; however, future studies should validate this correlation using biophysical and biochemical assays that directly measure ligand affinity or potency (SPR, enzymatic assays), which were not the focus of this study.

### Posaconazole rescues TDP-43 pathology in human iPSC-derived motor neurons treated with low dose sodium arsenite

Having demonstrated that posaconazole is a viable alternative to ketoconazole due to its known safer profile and enhanced effects on TDP-43, we wanted to demonstrate its ability to reduce TDP-43 pathology in a more physiologically relevant cellular model of ALS. Thus, we differentiated the human KOLF2.1J iPSC line into motor neurons characterized by strong nuclear TDP-43 expression, positive staining for STMN2 and extensive neuronal networks, as shown in the immunohistochemistry images in Fig. 6A-B.

**Fig. 6.**
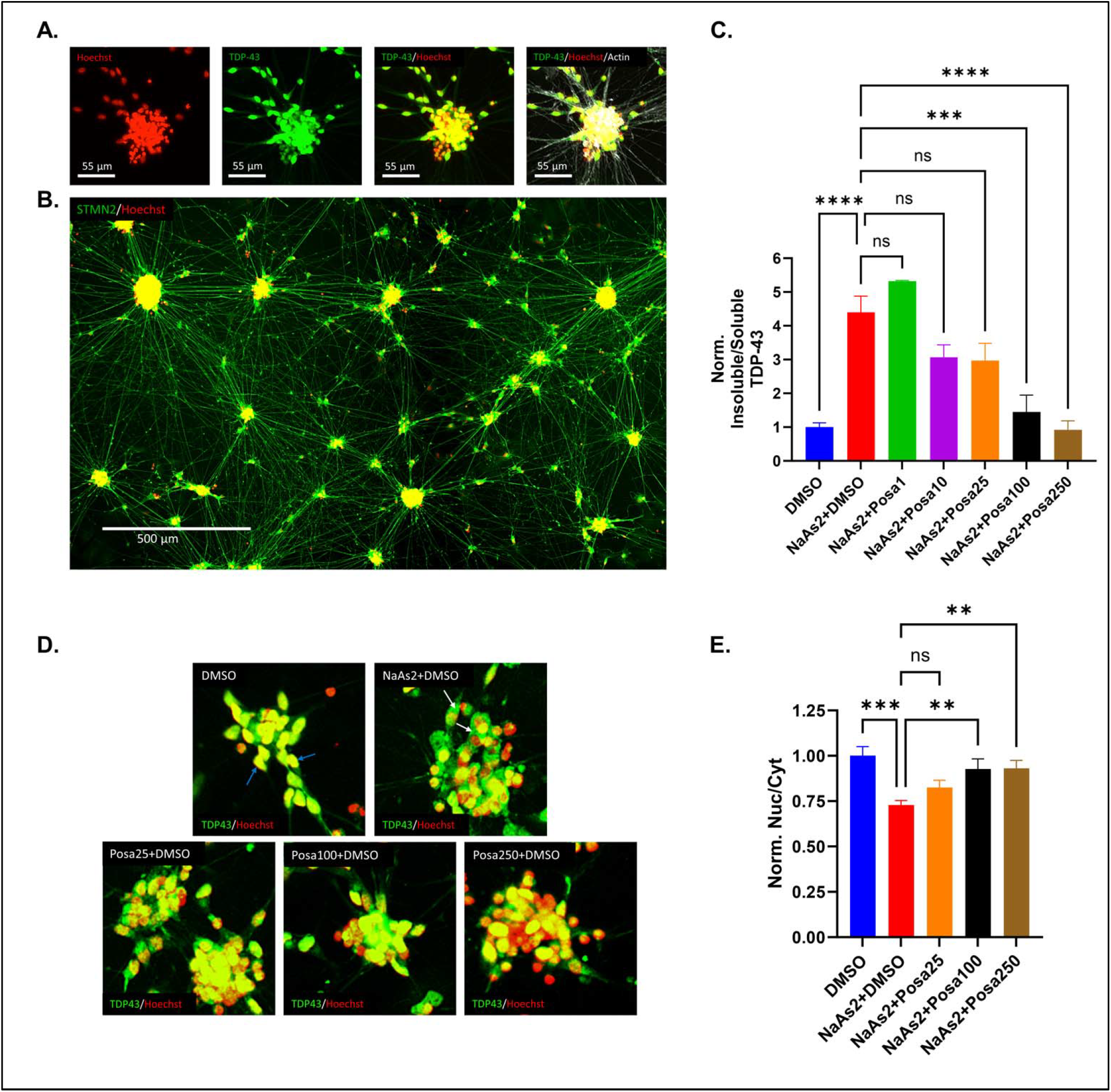
Posaconazole rescues TDP-43 pathology induced by low dose NaAs2 in human iPSC-derived motor neurons. **(A)** Nuclear TDP-43 expression pattern in motor neuron structures. **(B)** STMN2 expression patterns and large motor neuron network structures. **(C)** Biochemical characterization of insoluble-to-soluble ratio of TDP-43 in DMSO, NaAs2+DMSO and NaAs2+posaconazole (at 1, 10, 25, 100 and 250 nM) treated motor neurons. Data shown in (C) are mean ± SEM of N=3-9 independent treatments, analyzed via a one-way ANOVA with Bonferroni correction for multiple comparisons relative to DMSO/NaAs2 treated cells (***p < 0.001, ****p<0.0001). **(D)** Images of motor neuron cluster with +/-10 μM NaAs2 co-treated with DMSO or posaconazole at 25, 100 and 250 nM. **(E)** Normalized nuclear-to-cytoplasmic ratios (Nuc/Cyt) for the different treatments. Data shown in (D-E) are mean ± SEM of N=3 independent treatments (with a total of 4-12 technical replicates/images), analyzed via a one-way ANOVA with Bonferroni correction for multiple comparisons relative to DMSO/NaAs2 treated cells (*p < 0.05, **p < 0.01, ***p < 0.001, ****p<0.0001).

Using these motor neuron populations, we induced TDP-43 pathology by low dose sodium arsenite (NaAs2) (10 μM) for 24 hours, which caused significant TDP-43 pathology as quantified via western blotting as the ratio of insoluble/soluble TDP-43 (Fig. 6C, Fig. S12-13). Low dose NaAs2 is known to induce oxidative stress by generating reactive oxygen species (ROS), known to trigger TDP-43 cytoplasmic mislocalization and aggregation (*3, 42, 43*). Pre-treatment with posaconazole for 24 hours prior to NaAs2 treatment was able to significantly inhibit the increase in insoluble/soluble TDP-43 ratio in a dose dependent manner, reaching untreated levels at the dose of 250 nM (Fig. 6C). As shown in Fig. S13, the decrease in the insoluble/soluble ratio induced by posaconazole under NaAs2 treatment is attributed to an increase in soluble TDP-43, concomitant with a decrease in insoluble TDP-43. Finally, we quantified TDP-43 cytoplasmic mislocalization under NaAs2 treatment, which resulted in a 25% decrease in the Nuc/Cyt ratio of the protein relative untreated motor neurons (Fig. 6D-E). Pre-treatment with posaconazole at 100 and 250 nM were able to significantly rescue this deficit, in agreement with the western blot results (Fig. 6E).

## Discussion

Multiple recent studies point to sterol dysregulation as a key player in ALS (*44-47*). Most studies have focused on peripheral cholesterol levels, mostly agreeing there is an overall increase in serum and CSF cholesterol in ALS patients relative to healthy controls (*44, 48*). It is still unclear whether the elevated peripheral cholesterol levels are detrimental or protective (mixed reports supporting both hypothesis), and comprehensive connecting CNS cholesterol levels to disease progression or severity is still lacking (*44*). Interestingly, a recent study determined that cholesterol levels are significantly increased in skeletal muscle cells of ALS patients relative to controls, and that the degree of cholesterol accumulation correlates with disease severity (*49*). Cholesterol esters and triacylglycerol levels are elevated in the spinal cords of sporadic ALS patients, and byproducts of these species can be toxic to motor neurons (*47*). This agrees with our observation of SREBP2 dysregulation in the cervical and lumbar spinal cord regions of ALS patients, as SREBP2 is downregulated when intracellular cholesterol levels are increased. Longitudinal studies of cholesterol and its related molecules’ levels in plasma, serum, CSF and skeletal muscle biopsies of ALS patients throughout disease progression will be crucial to understand the full contribution of this pathway to ALS. In addition, characterization of cholesterol and related metabolites in ALS-relevant tissues such as the spinal cord in post-mortem samples will give a more complete picture.

Given the implication of cholesterol metabolism in ALS, the usage of statins is being actively explored as a potential therapeutic (*50, 51*). A recent pharmacoepidemiologic study identified that the use of cholesterol-lowering statins that target HMG-CoA reductase (lovastatin, simvastatin, atorvastatin, rosuvastatin and pravastatin) significantly reduced the risk of developing ALS in individuals aged 66-90 residing in the United States (*50*). In support of these clinical results, it was recently shown that statins can induce an increase in STMN2 levels (a critical protein for axonal integrity in the context of ALS (*52*)) in SH-SY5Y cells (*53*). In addition, a recent pre-clinical study using poly-GA C9orf72 mice showed that dysregulated cholesterol metabolism can be corrected with cholesterol depletion via 2-hydroxypropyl-β-cyclodextrin (a cholesterol-sequestering molecule), which led to prolonged lifespan relative to untreated mice (*54*). Altogether, these pre-clinical and clinical results implicate cholesterol depletion as a potential therapeutic strategy for ALS. Our results further support these clinical findings and shed light on a promising mechanism connecting CYP51 inhibition, cholesterol depletion and rescue of TDP-43 pathology that could be leveraged as a therapeutic for ALS. Posaconazole may be a good candidate as a repurposed ALS therapeutic for several reasons. First, it was able to fully restore downregulated SREBP2 levels, which are known to be affected by TDP-43 pathology in oligodendrocytes, disrupting the expression patterns of many cholesterol pathway associated genes and myelin production crucial for their supporting role in the CNS (*9*). As shown here, SREBP2 expression levels are also reduced in cervical and lumbar spinal cord tissue samples from ALS/FTD patients, although whether this decrease is due to TDP-43 pathology remains unclear and should be further investigated. By reducing cholesterol levels and increasing SREBP2 expression, posaconazole may be able to re-balance the cholesterol synthesis pathway in ALS/FTD patients.

Second, posaconazole rescued TDP-43 mislocalization and aggregation by activating autophagy and increasing the expression of crucial heat shock proteins for proteostasis such as HSPB5, HSPB8, DNAJB2 and HSPB1. HSPB5 (also known as αB-crystallin) prevents the aggregation of multiple protein targets including the Parkinson’s disease-related alpha-synuclein and ALS-related SOD1 (*55, 56*). In addition, upregulation of the HSPB5 ortholog in Drosophila has been shown to reduce TDP-43 neurotoxicity (*57*). Interestingly, as shown by Hu et al., HSPB5 was the most upregulated target in a heat shock protein panel following lanosterol treatment (*35*). Coincidentally, HSPB5 was the target with highest fold increase for both ketoconazole and posaconazole in this study, and these compounds are known to trigger lanosterol accumulation by inhibiting CYP51 (*33, 58*). HSPB8 has been shown to reduce mislocalization and accumulation of neurotoxic TDP-43 fragments in immortalized motor neurons (*59*). DNAJB2 was identified as a potent anti-aggregation TDP-43 chaperone (*37*). HSPB1 has been recently shown to inhibit the de-mixing of TDP-43 into amyloid fibrils and is significantly reduced in surviving motor neurons in spinal cord slices of ALS patients relative to healthy controls (*38*). Moreover, the same study identified HSPB1 genetic variants with association to ALS (*38*). Interestingly, HSPB1 was the heat shock protein most downregulated in this study due to TDP-43 overexpression, which was rescued by posaconazole treatment almost fully back to healthy control cells. Finally, posaconazole upregulated the levels of HSP90A and HSP90B, which also modulate TDP-43 misfolding and aggregation (*60*). Both ketoconazole and posaconazole failed to increase HSP70 levels at 24-hours; however, we speculate that given the tight regulation of HSP70 in the heat shock pathway along with the efficient clearance of TDP-43 by these molecules, this observation may reflect the resolution of the heat shock response. Indeed, the therapeutic compound arimoclomol, which reduces the levels of toxic lysosomal unesterified cholesterol in the context of Niemman-pick type C disease, is able to induce a rapid increase in HSP70 at early timepoints, which is then resolved at later timepoints (*61, 62*). Alternatively, in the case that these compounds are unable to activate HSP70, the significant redundancy of the heat shock protein pathway (*63, 64*), and their ability to activate autophagy still explains why these compounds were effective at reducing insoluble TDP-43. Regarding autophagy, which is severely impaired in ALS (*65*), several activators of this pathway such as rapamycin, tamoxifen, monepantel and bosutinib are currently being evaluated for ALS in Phase I and/or II clinical trials (*66*).

Third, posaconazole is prescribed as an oral medication with lower hepatotoxicity than ketoconazole (*67*) (*68*), and it is used to treat CNS fungal infections with encouraging success rates, for which higher concentrations and longer-term treatments relative to topical fungal infections are required, but well-tolerated (*69*). Here, we show that posaconazole was able to block TDP-43 pathology induced by NaAs2 oxidative stress in what is arguably one of the most relevant CNS cell types in the context of ALS, that is, motor neurons. In agreement with our results in immortalized human cells, posaconazole inhibited the increase in TDP-43 pathology in terms of insoluble-to-soluble ratio and mislocalization. Given its effects at 100 and 250 nM, we hypothesize that if posaconazole is able to penetrate the CNS at high enough concentrations, it would have a beneficial effect by restoring SREBP2 expression and lowering TDP-43 burden.

In conclusion, we have demonstrated that the known anti-fungal azole-based inhibitors ketoconazole and posaconazole have strong activity towards TDP-43 pathology in immortalized human cells, that their effects are correlated with their predicted affinities towards human CYP51, and that they act in a cholesterol-dependent mechanism. From these experiments, we were able to identify posaconazole, a more efficacious and safer alternative to ketoconazole, which rescued TDP-43 pathology in human motor neurons. Altogether, these results support further pursuing posaconazole and other cholesterol pathway-modulating compounds as potential ALS/FTD treatments.

## Methods

### Multiple sequence alignment and structural alignment to human CYP51

CYP51 sequences for H. sapiens, A. fumigatus, T. cruzi, S. parasitica and C. albicans were obtained from NCBI’s Gene database using the following NCBI Sequence Reference numbers: NP_000777.1, AAP33132.1, XP_012204080.1, UTE79867.1, KAH1581877.1. The sequences were aligned using multiple sequence alignment via ClustalW (*70*), from which global sequence identities were calculated. Known motif sequences (substrate recognition sites SRS1-6, oxygen binding motif, heme binding motif, EXXR motif and PXR motif) were extracted from the multiple sequence alignment and used to calculate conserved motif sequence identities shown in Fig. S2. Structural alignment of Boltz-2 predicted structures for H. sapiens and the 4 fungal CYP51 sequences against PDB ID: 3LD6 (H. sapiens) were conducted using PyMOL.

### Cell culture

HEK293T cells (ATCC) were maintained in full Gibco DMEM media (10% FBS, 1% Pen/Strep, 1% GlutaMax) at 37 °C, 5% CO2, and humidity control. All transfections were carried out using Thermo Fisher Lipofectamine 3000 at a DNA:P3000:L3000 ratio of 1:2:3 (μg DNA: μL P3000: μL L3000), with 2.5 μg DNA per 1e6 seeded cells. For cholesterol quantification, cells were incubated in cholesterol-free OptiMEM media.

### TDP-43 mislocalization and aggregation imaging assay

HEK293T cells were seeded at 0.16e6 cells per well in 24-well plates (Eppendorf). After 24 hours of incubation, each well was transfected with 0.08 μg TDP-43-Ng supplemented with 0.32 μg empty vector for a total of 0.4 ug DNA per well (2.5 μg DNA/1e6 cells). After 24 hours of incubation, cells were pre-treated with 0.2% DMSO control and 10 μM inhibitors for 1.5 hours. Next, Hoechst stain was added at a 1:50k dilution and the cells were incubated for an additional 30 minutes. Images of 2-hour treatments shown in Fig. 1 were collected at this stage. Next, ultra-pure water control or 0.1 M sorbitol were added to wells. Plates were imaged for nuclear, TDP-43-mNg and brightfield channels after 24 hours after addition of drug in a Molecular Devices Pico ImageXpress microscope. The same procedure was followed for dose responses of ketoconazole, posaconazole.

Images were analyzed using MetaXpress High Content Image Analysis software with two custom analysis workflows that quantify number of cytoplasmic puncta per cell and nuclear/cytoplasmic localization based on intensity thresholding of the green fluorescence channel and masking using the Hoechst channel.

### Soluble and insoluble protein fraction extractions

Soluble and insoluble protein fractions were extracted using the protocol established by Hans et. al. as reference (*22*). Briefly, TDP-43-mNg-expressing cells used for fluorescence imaging under DMSO, DMSO/sorbitol, ketoconazole/sorbitol and posaconazole/sorbitol were harvested and washed in PBS prior to lysis in RIPA buffer. After 5 minutes of lysis on ice, the samples were spun down at 14,000 g for 15 minutes. Bicinchoninic acid assays (BCA, Thermo Scientific) were conducted on the supernatant to quantify total protein content. The supernatant was saved as the soluble fraction, and the remaining pellet (insoluble fraction) was solubilized with 8 M urea solution for 1 hour at 60 °C, and overnight at RT in an orbital shaker. Both fractions were prepared in 1X Laemmli buffer supplemented with 20 μM DTT.

### Immunoblotting

SDS-PAGE gels were run at 120V using 12% BioRad gels with 10-20 μg of protein loaded in each well. Gels were transferred into nitrocellulose membranes using the BioRad TurboTransfer protocol and stained for total protein using Ponceau S. Finally, blots were probed with TDP-43 (Proteintech 10782-2-AP), LC3A/B (Cell Signaling D3U4C), alpha-tubulin (Proteintech HRP-66031) and histone 3 (Invitrogen MA5-45047) at 1:1000-1:5000 dilution overnight at 4 °C and detected using HRP-conjugated secondary antibody at 1:10000 dilution imaged with SuperSignal West Pico PLUS chemiluminescent substrate (Thermo Scientific).

### Cholesterol measurement under TDP-43 overexpression conditions

HEK293T cells were plated in 12-wells, incubated for 24 hours and transfected with unlabeled TDP-43 at a ratio of DNA/cells of 2.5 μg/1e6 cells in cholesterol-depleted conditions (Opti-MEM) to rule out cholesterol uptake from full media. Two hours after transfection, cells were treated with DMSO, 10 μM ketoconazole and 10 μM posaconazole for an additional 24 hours. Cells were harvested, of which 10% were used for total protein quantification via BCA. The remaining cells were pelleted, resuspended in a 7:11:0.1 mixture of chloroform:isopropanol:IGEPAL CA-630 and incubated with shaking for 10 minutes at 1500 rpm to extract lipids. Next, the extractions were pelleted for 10 minutes at 13,000 g, and the supernatants were left to air dry overnight. The dried lipids were quantified for free cholesterol using the Amplex Red Cholesterol Assay Kit (Thermo Fisher), following manufacturers protocol. Cholesterol concentrations were normalized to total protein concentrations and to untreated cell levels.

### Analysis of NYGC ALS Consortium dataset for SREBP2 expression across tissue types in ALS vs. controls

The NYGC ALS Consortium bulk RNA-sequencing dataset by tissue type and associated metadata was downloaded from Zenodo (https://doi.org/10.5281/zenodo.20090702). Transcript per million (TPM) counts for each tissue were transformed using a log2(TPM+0.01) transformation, grouping the data by controls or ALS/FTD samples.

### RT-qPCR

HEK293T cells were seeded at 0.8e6 cells per well in 6-well plates and incubated for 24 hours. After incubation, cells were transfected with 2 μg of unlabeled full-length TDP-43 (2.5 μg DNA/1e6 cells). Two hours after transfection, cells were treated with 10 μM ketoconazole and posaconazole (as well as 0.2% matching DMSO control) and incubated for an additional 24 hours.

Total RNA for each condition was isolated using Qiagen RNAeasy extraction protocol. After quality validation via 280/260 measurements, total RNA was reversed transcribed using Thermo Fisher High-Capacity RNA-to-cDNA kit. PCR reactions probing for endogenous GAPDH (housekeeping) and SREBP2 were prepared following the Thermo Fisher SYBR Green qPCR protocol and using the following specific primers (IDT) for endogenous mRNA (designed using NCBI’s primer designing tool):

GAPDH

Forward: 5’-AATGGGCAGCCGTTAGGAAA-3’

Reverse: 5’-GCGCCCAATACGACCAAATC-3’

SREBP2:

Forward: 5’-TGTGTCCTCACCTTCCTGTGCCT-3’

Reverse: 5’-TCCAGTCAAACCAGCCCCCAGA-3’

HSP90A:

Forward: 5’-TCTGCCTCTGGTGATGAGATGG-3’

Reverse: 5’-CGTTCCACAAAGGCTGAGTTAGC-3’

HSP90B:

Forward: 5’-CTCTGTCAGAGTATGTTTCTCGC-3’

Reverse: 5’-GTTTCCGCACTCGCTCCACAAA-3’

HSP70:

Forward: 5’-ACCTTCGACGTGTCCATCCTGA-3’

Reverse: 5’-TCCTCCACGAAGTGGTTCACCA-3’

HSPB5:

Forward: 5’-ACTTCCCTGAGTCCCTTCTACC-3’

Reverse: 5’-GGAGAAGTGCTTCACATCCAGG-3’

HSPB1:

Forward: 5’-CTGACGGTCAAGACCAAGGATG-3’

Reverse: 5’-GTGTATTTCCGCGTGAAGCACC-3’

DNAJB2:

Forward: 5’-GGAGATTTACGACCGCTATGGC-3’

Reverse: 5’-GCAAAAGGGTCTCCACTCCCAA-3’

HSPB8:

Forward: 5’-CAGAGGAGTTGATGGTGAAGACC-3’

Reverse: 5’-ACTGTCACAGGATCCACCTCTG-3’

RT-qPCR reactions were conducted using a Roche LightCycler 96 plate reader and analyzed using Roche’s PCR analysis software. Expression was quantified using the *2*^-ΔΔCt^ method relative to untreated/untransfected HEK293T cells (*71*).

### Production of MBCD:cholesterol complexes for cholesterol supplementation experiments

MBCD and cholesterol were dissolved in ultra-pure filtered water at a 6:1 molar ratio (final concentrations of 6 mM and 1 mM) with constant heating and stirring at 50-60 °C for 1 hour or until the solution became opalescent. The solution was briefly sonicated (3 × 15 s pulses at 40% amplitude) and immediately filtered through a 22 μM filter to remove large particulate. Complexes were stored in glass vials at 4 °C for up to a week until use. All cholesterol supplementation treatments were performed at a final supplemented cholesterol concentration of 100 μM.

### Molecular docking of azole inhibitors to human CYP51

The ketoconazole-bound human CYP51 structure (PDB ID: 3LD6) was imported into Maestro (Schrödinger) and processed in order to add missing hydrogens and residues. Next, ketoconazole was used to define the binding site using the Receptor Grid Generation module in Glide (*72*). In order to enforce azole-heme interactions for all inhibitors, a metal coordination positional constraint was generated. Using the Ligand Docking module, the 8 inhibitors and DMSO were docked CYP51. Docking was performed with standard precision (SP), allowing flexibility for ligands. A short 200 step minimization was conducted post-docking in Maestro. Docking scores were correlated to fungal CYP51 inhibition data, and mislocalization/aggregation data in GraphPad Prism 9.

Boltz-2 binding affinity predictions were conducted using template scripts provided by the authors (*41*) and implemented using the following FASTA sequence for human CYP51 (extracted from NCBI) and SMILES representations for heme and the 8 inhibitors (extracted from PubChem):

CYP51 FASTA: MAKKTSSKGKLPPYIFSPIPFLGHAIAFGKSPIEFLENAYEKYGPVFSFTMVGKTFTYLLGSDAAALLFNSKNEDLNAEDVYSRLTTPVFGKGVAYDVPNPVFLEQKKMLKSGLNIAHFKQHVSIIEKETKEYFESWGESGEKNVFEALSELIILTASHCLHGKEIRSQLNEKVAQLYADLAGGFSHAAWLLPGWLPLPSFRRRDRAHREIKDIFYKAIQKRRQSQEKIDDILQTLLDATYKDGRPLTDDEVAGMLIGLLLAGQATSSTTSAWMGFFLARDKTLQKKCYLEQKTVCGENLPPLTYDQLKDLNLLDRCIKETLRLRPPIMIMMRMARTPQTVAGYTIPPGHQVCVSPTVNQRLKDSWVERLDFNPDRYLQDNPASGEKFAYVPFGAGRHRCIGENFAYVQIKTIWSTMLRLYEFDLIDGYFPTVNYTTMIHTPENPVIRYKRRS

Heme SMILES:

CC1=C(C2=CC3=NC(=CC4=C(C(=C([N-]4)C=C5C(=C(C(=N5)C=C1[N-

]2)C=C)C)C=C)C)C(=C3CCC(=O)[O-])C)CCC(=O)[O-].[Fe]

Ketoconazole SMILES: CC(=O)N1CCN(CC1)C2=CC=C(C=C2)OC[C@@H]3CO[C@@](O3)(CN4C=CN=C4)C5=C(C=C(C=C5)Cl)Cl

Posaconazole SMILES: CC[C@@H]([C@H](C)O)N1C(=O)N(C=N1)C2=CC=C(C=C2)N3CCN(CC3)C4=CC=C(C=C4)OC[C@H]5C[C @](OC5)(CN6C=NC=N6)C7=C(C=C(C=C7)F)F

Fluconazole SMILES:

C1=CC(=C(C=C1F)F)C(CN2C=NC=N2)(CN3C=NC=N3)O

Voriconazole SMILES:

C[C@@H](C1=NC=NC=C1F)[C@](CN2C=NC=N2)(C3=C(C=C(C=C3)F)F)O

Clotrimazole SMILES:

C1=CC=C(C=C1)C(C2=CC=CC=C2)(C3=CC=CC=C3Cl)N4C=CN=C4

Fenticonazole SMILES:

C1=CC=C(C=C1)SC2=CC=C(C=C2)COC(CN3C=CN=C3)C4=C(C=C(C=C4)Cl)Cl

Cyproconazole SMILES:

CC(C1CC1)C(CN2C=NC=N2)(C3=CC=C(C=C3)Cl)O

Itraconazole SMILES:

CCC(C)N1C(=O)N(C=N1)C2=CC=C(C=C2)N3CCN(CC3)C4=CC=C(C=C4)OC[C@H]5CO[C@](O5)(CN6C= NC=N6)C7=C(C=C(C=C7)Cl)Cl

### Differentiation of human iPSCs into motor neurons and low dose sodium arsenite model

Motor neurons were generated using an accelerated version of the Cedars Sinai differentiation protocol using the KOLF2.1J hiPSC line (*73-75*), maintained in Essential 8 (E8) media. Briefly, hiPSCs were seeded at 10,000 cell/cm^2^ in Geltrex coated plates for 24 hours and differentiated using 5 distinct media preparations across 7 days. On day 1, cells were plated in day 1 media consisting of Essential 6 (E6) media supplemented with 100 nM retinoic acid, 500 nM LDN-193189 and 100 nM BGJ398. After 24 hours, media was replaced with E6 supplemented with 100 nM retinoic acid, 4 μM CHIR99021, 20 ng/mL FGF2 and 500 nM A8301. After 24 hours, media was replaced with E6 supplemented with 100 nM retinoic acid, 100 nM BGJ398, 250 nM wntC59 and 1000 nM SAG. After 24 hours, media was replaced with E6 supplemented with 100 nM retinoic acid, 20 ng/mL FGF2, 500 nM A8301 and 1000 nM SAG. After 24 hours, this same media (day 4) was replenished for an additional 24 hours. At day 6, cells were fed for 48 hours with S3 media, consisting of equal parts IMDM and Ham’s F12 media supplemented with 1x B27, 1x N2, 1x NEAA, 1x Anti-anti, 500 nM retinoic acid, 100 nM SAG, 100 nM Compound E, 2.5 μM DAPT, 100 nM db-cAMP, 100 μM ascorbic acid, 10 ng/mL BDNF and 10 ng/mL GDNF. At day 8, precursor motor neurons were harvested using Accutase and frozen down with CryoStor CS10 in vials containing 4.5-9e6 cells in liquid nitrogen.

For NaAs2 treatments, single precursor motor neuron vials were thawed and plated in Geltrex-coated 6-well plates in S3 media (see Table S3) with ROCK inhibitor at a density of 20,000 cells/cm^2^ for terminal motor neuron differentiation. After 24 hours, ROCK inhibitor media was removed, and cells were differentiated for an additional 6 days in S3 media with daily media switching. After differentiation, motor neurons were pre-treated with DMSO, 1, 10, 25, 100 and 250 nM posaconazole (0.1% DMSO final concentration) for 24 hours. Next, NaAs2 was added to each well at a final concentration of 10 μM and incubated for an additional 24 hours before processing for insoluble and soluble protein fraction extractions via western blotting, as described in the previous sections.

### Immunohistochemical imaging of motor neurons

Motor neurons were differentiated in Geltrex-coated 96-well plates as described in the previous section. After full differentiation, the cells were prepared for immunohistochemical analysis. S3 media was removed, followed by a PBS wash and 15-minute fixing in room temperature 4% paraformaldehyde (ChemCruz) in PBS. Next, PFA was removed, followed by a PBS wash and a 10-minute incubation in permeabilizing solution (0.1% Triton-X in PBS). Next, cells were washed and blocked in 5% BSA in PBS for 1 hour, followed by incubation with primary antibodies (TDP-43 and STMN2 (Invitrogen 720178), 1:1000 dilution in 5% BSA in PBS) overnight at 4 °C. Next, cells were washed with PBS and incubated with AlexaFluor 488 tagged secondary antibody for 2 hours at room temperature. Finally, cells were washed in PBS, incubated with phalloidin rhodamine actin stain and Hoechst nuclear stain for 30 minutes, followed by a final PBS wash prior to imaging in a Molecular Devices Pico ImageXpress microscope.

### Statistical analysis

Multiple comparisons were conducted via ordinary one-way or two-way ANOVAs with post-hoc Bonferroni corrections for multiple comparisons. For SREBP2 expression across tissue types, multiple comparisons were conducted via a two-way ANOVA with Benjamini-Hochberg False Discovery Rate correction for multiple comparisons. All statistical analysis was done in GraphPad Prism 9.

## Author contributions

Conceptualization: NNK, JNS

Methodology: NNK, SZ, AR, NS, NV, EEL, ARB

Investigation: NNK, SZ, AR, NS, AR, NV, EEL, ARB

Visualization: NNK

Supervision: JRD, ARB, JNS

Writing—original draft: NNK

Writing—review & editing: NNK, ARB, JNS

## Data and materials availability

All data needed to evaluate the conclusions are present in the paper and/or the Supplementary Materials. The datasets, scripts and materials generated in this study are available from the corresponding author on reasonable request.

## Supplementary Materials

The supplementary materials include additional imaging, molecular modeling, sequence analysis and biochemical experimental data.

## References

1. J. Zeng, C. Luo, Y. Jiang, T. Hu, B. Lin, Y. Xie, J. Lan, J. Miao, Decoding TDP-43: the molecular chameleon of neurodegenerative diseases. Acta Neuropathologica Communications 12, 205 (2024).

2. C. Scialò, W. Zhong, S. Jagannath, O. Wilkins, D. Caredio, M. Hruska-Plochan, F. Lurati, M. Peter, E. De Cecco, L. Celauro, A. Aguzzi, G. Legname, P. Fratta, M. Polymenidou, Seeded aggregation of TDP-43 induces its loss of function and reveals early pathological signatures. Neuron 113, 1614-1628.e1611 (2025).

3. K. Oiwa, S. Watanabe, K. Onodera, Y. Iguchi, Y. Kinoshita, O. Komine, A. Sobue, Y. Okada, M. Katsuno, K. Yamanaka, Monomerization of TDP-43 is a key determinant for inducing TDP-43 pathology in amyotrophic lateral sclerosis. Science Advances 9, (2023).

4. S.-C. Ling, M. Polymenidou, Don W. Cleveland, Converging Mechanisms in ALS and FTD: Disrupted RNA and Protein Homeostasis. Neuron 79, 416–438 (2013).

5. N. Nathan Kochen, M. Murray, S. Zafari, N. Vunnam, E. E. Liao, L. Chen, A. R. Braun, J. N. Sachs, Fluorescence Lifetime-Based FRET Biosensors for Monitoring N Terminal Domain-Dependent Interactions of TDP-43 in Living Cells: A Novel Approach for ALS and FTD Drug Discovery. ACS Chemical Neuroscience, (2025).

6. T. Y. Hargrove, L. Friggeri, Z. Wawrzak, A. Qi, W. J. Hoekstra, R. J. Schotzinger, J. D. York, F. P. Guengerich, G. Lepesheva, Structural analyses of <em>Candida albicans</em> sterol 14α-demethylase complexed with azole drugs address the molecular basis of azole-mediated inhibition of fungal sterol biosynthesis. Journal of Biological Chemistry 292, 6728–6743 (2017).

7. N. Strushkevich, S. A. Usanov, H.-W. Park, Structural Basis of Human CYP51 Inhibition by Antifungal Azoles. Journal of Molecular Biology 397, 1067–1078 (2010).

8. N. Egawa, Y. Izumi, H. Suzuki, I. Tsuge, K. Fujita, H. Shimano, K. Izumikawa, N. Takahashi, K. Tsukita, T. Enami, M. Nakamura, A. Watanabe, M. Naitoh, S. Suzuki, T. Seki, K. Kobayashi, T. Toda, R. Kaji, R. Takahashi, H. Inoue, TDP-43 regulates cholesterol biosynthesis by inhibiting sterol regulatory element-binding protein 2. Scientific Reports 12, 7988 (2022).

9. W. Y. Ho, J. C. Chang, K. Lim, A. Cazenave-Gassiot, A. T. Nguyen, J. C. Foo, S. Muralidharan, A. Viera-Ortiz, S. J. M. Ong, J. H. Hor, I. Agrawal, S. Hoon, O. A. Arogundade, M. J. Rodriguez, S. M. Lim, S. H. Kim, J. Ravits, S. Y. Ng, M. R. Wenk, E. B. Lee, G. Tucker-Kellogg, S. C. Ling, TDP-43 mediates SREBF2-regulated gene expression required for oligodendrocyte myelination. J. Cell Bio. 220, (2021).

10. J. A. Barnes-Vélez, F. B. Aksoy Yasar, J. Hu, Myelin lipid metabolism and its role in myelination and myelin maintenance. The Innovation 4, (2023).

11. X. A.-O. Zhou, S. A.-O. Shin, C. He, Q. Zhang, M. A.-O. Rasband, J. Ren, C. Dai, R. I. Zorrilla-Veloz, T. Shingu, L. Yuan, Y. Wang, Y. Chen, F. Lan, J. A.-O. Hu, Qki regulates myelinogenesis through Srebp2-dependent cholesterol biosynthesis. LID - 10.7554/eLife.60467 [doi] LID - e60467.

12. S. M. Colgan, D. Tang, G. H. Werstuck, R. C. Austin, Endoplasmic reticulum stress causes the activation of sterol regulatory element binding protein-2. The International Journal of Biochemistry & Cell Biology 39, 1843–1851 (2007).

13. N. Q. Wang, P. X. Sun, Q. Q. Shen, M. Y. Deng, Cholesterol Metabolism in CNS Diseases: The Potential of SREBP2 and LXR as Therapeutic Targets. Molecular Neurobiology, (2025).

14. K. E. Shapira, G. Shapira, E. Schmukler, M. Pasmanik-Chor, N. Shomron, R. Pinkas-Kramarski, Y. I. Henis, M. Ehrlich, Autophagy is induced and modulated by cholesterol depletion through transcription of autophagy-related genes and attenuation of flux. Cell Death Discovery 7, 320 (2021).

15. J. Cheng, Y. Ohsaki, K. Tauchi-Sato, A. Fujita, T. Fujimoto, Cholesterol depletion induces autophagy. Biochemical and Biophysical Research Communications 351, 246–252 (2006).

16. R. Aron, B. Pandya, D. Tardiff, J. Piotrowski, M. Lucas, B. Le Bourdonnec, K. Rhodes, R. Scannevin, W. I. P. Organization, Ed. (Yumanity Therapeutics, Inc, 2019), chap. WO 2019/246494.

17. N. P. Taylor, in Fierce Biotech. (Online, 2022).

18. H. K. Greenblatt, D. J. Greenblatt, Liver injury associated with ketoconazole: Review of the published evidence. The Journal of Clinical Pharmacology 54, 1321–1329 (2014).

19. T. Y. Hargrove, L. Friggeri, Z. Wawrzak, S. Sivakumaran, E. M. Yazlovitskaya, S. W. Hiebert, F. P. Guengerich, M. R. Waterman, G. I. Lepesheva, Human sterol 14α-demethylase as a target for anticancer chemotherapy: towards structure-aided drug design.

20. A. G. Warrilow, C. M. Hull, N. J. Rolley, J. E. Parker, W. D. Nes, S. N. Smith, D. E. Kelly, S. L. Kelly, Clotrimazole as a potent agent for treating the oomycete fish pathogen Saprolegnia parasitica through inhibition of sterol 14α-demethylase (CYP51).

21. B. Celia-Sanchez, B. Mangum, M. Brewer, M. Momany, Analysis of Cyp51 protein sequences shows 4 major Cyp51 gene family groups across fungi. G3: Genes, Genomes, Genetics, (2022).

22. F. Hans, H. Glasebach, P. A.-O. Kahle, Multiple distinct pathways lead to hyperubiquitylated insoluble TDP-43 protein independent of its translocation into stress granules.

23. C. M. Dewey, C. B. C. F. Sephton, D. R. Dries, P. r. Mayer, S. K. Good, B. A. Johnson, J. Herz, G. Yu, TDP-43 is directed to stress granules by sorbitol, a novel physiological osmotic and oxidative stressor. Mol. Cell Bio. 31, 1098–1108 (2011).

24. G. I. Lepesheva, L. Friggeri, M. R. Waterman, CYP51 as drug targets for fungi and protozoan parasites: past, present and future. Parasitology, (2018).

25. J. Zhu, K. Mounzih, E. F. Chehab, N. Mitro, E. Saez, F. F. Chehab, Effects of FoxO4 overexpression on cholesterol biosynthesis, triacylglycerol accumulation, and glucose uptake. Journal of Lipid Research 51, 1312–1324 (2010).

26. R. A. DeBose-Boyd, Feedback regulation of cholesterol synthesis: sterol-accelerated ubiquitination and degradation of HMG CoA reductase. Nature Cell Research 18, 609–621 (2008).

27. J. L. Goldstein, R. A. DeBose-Boyd, M. S. Brown, Protein Sensors for Membrane Sterols. Cell 124, 35–46 (2006).

28. J. Humphrey, S. Venkatesh, R. Hasan, J. T. Herb, K. de Paiva Lopes, F. Küçükali, M. Byrska-Bishop, U. S. Evani, G. Narzisi, D. Fagegaltier, K. Sleegers, H. Phatnani, D. A. Knowles, P. Fratta, T. Raj, Integrative transcriptomic analysis of the amyotrophic lateral sclerosis spinal cord implicates glial activation and suggests new risk genes. Nature Neuroscience, (2023).

29. J. Humphrey, A. Oku, M. Byrska-Bishop, A. O. Basile, U. S. Evani, A. Corvelo, A. Tokolyi, K. Bp, A. Réal, Y. Kim, M. L. Bond, W. E. Clarke, R. Fu, H. Geiger, S. Chang, T. Naito, B. Jang, R. Musunuri, W. H. Dredge, R. Al-Abri, B. N. Hoover, D. Manaa, J. McClintock, F. P. Singh, M. H. Pedersen, A. Runnels, N. Propp, S. Fennessey, H.-H. Won, M. C. Zody, G. Narzisi, N. Robine, T. Lappalainen, D. Fagegaltier, G. Gürsoy, D. A. Knowles, T. Raj, N. A. Consortium, M. B. Harms, H. Phatnani, The New York Genome Center ALS Consortium resource integrates postmortem tissue transcriptomics and whole genome sequencing to empower biological discovery. medRxiv, 2026.2004.2029.26350889 (2026).

30. A. M. G. Ragagnin, S. Shadfar, M. Vidal, M. S. Jamali, J. D. Atkin, Motor Neuron Susceptibility in ALS/FTD. Frontiers in Neuroscience, (2019).

31. R. L. Barry, A. Torrado-Carvajal, J. E. Kirsch, G. E. Arabasz, D. S. Albrecht, Z. Alshelh, O. Pijanowski, A. J. Lewis, M. Keegan, B. Reynolds, P. C. Knight, E. J. Morrissey, M. L. Loggia, N. Atassi, J. M. Hooker, S. Babu, Selective atrophy of the cervical enlargement in whole spinal cord MRI of amyotrophic lateral sclerosis. NeuroImage: Clinical, (2022).

32. E. Ratti, K. Domoto-Reilly, C. Caso, A. Murphy, M. Brickhouse, D. Hochberg, N. Makris, M. E. Cudkowicz, B. C. Dickerson, Regional prefrontal cortical atrophy predicts specific cognitive-behavioral symptoms in ALS-FTD. Brain Imaging and Behavior, (2021).

33. H. J. Kempen, K. Van Son, L. H. Cohen, M. Griffioen, H. Verboom, L. Havekes, Effect of ketoconazole on cholesterol synthesis and on HMG-COA reductase and LDL-receptor activities in Hep G2 cells. Biochemical Pharmacology 36, 1245–1249 (1987).

34. A. Efeyan, W. C. Comb, D. M. Sabatini, Nutrient-sensing mechanisms and pathways. Nature, (2015).

35. L.-D. Hu, J. Wang, X.-J. Chen, Y.-B. Yan, Lanosterol modulates proteostasis via dissolving cytosolic sequestosomes/aggresome-like induced structures. Biochimica et Biophysica Acta (BBA) - Molecular Cell Research 1867, 118617 (2020).

36. L. Zhao, X.-J. Chen, J. Zhu, Y.-B. Xi, X. Yang, L.-D. Hu, H. Ouyang, S. H. Patel, X. Jin, D. Lin, F. Wu, K. Flagg, H. Cai, G. Li, G. Cao, Y. Lin, D. Chen, C. Wen, C. Chung, Y. Wang, A. Qiu, E. Yeh, W. Wang, X. Hu, S. Grob, R. Abagyan, Z. Su, H. C. Tjondro, X.-J. Zhao, H. Luo, R. Hou, J. Jefferson, P. Perry, W. Gao, I. Kozak, D. Granet, Y. Li, X. Sun, J. Wang, L. Zhang, Y. Liu, Y.-B. Yan, K. Zhang, Lanosterol reverses protein aggregation in cataracts. Nature 523, 607–611 (2015).

37. H. J. Chen, J. C. Mitchell, S. Novoselov, J. Miller, A. L. Nishimura, E. L. Scotter, C. A. Vance, M. E. Cheetham, C. E. Shaw, The heat shock response plays an important role in TDP-43 clearance: evidence for dysfunction in amyotrophic lateral sclerosis.

38. S. Lu, J. Hu, O. A. Arogundade, A. Goginashvili, S. Vazquez-Sanchez, J. K. Diedrich, J. Gu, J. Blum, S. Oung, Q. Ye, H. Yu, J. Ravits, C. Liu, J. R. Yates, D. W. Cleveland, Heat-shock chaperone HSPB1 regulates cytoplasmic TDP-43 phase separation and liquid-to-gel transition. Nature Cell Biology 24, 1378–1393 (2022).

39. M. Kadowaki, M. R. Karim, Cytosolic LC3 ratio as a quantitative index of macroautophagy.

40. Y. Watanabe, H. Tatebe, K. Taguchi, Y. Endo, T. Tokuda, T. Mizuno, M. Nakagawa, M. Tanaka, p62/SQSTM1-Dependent Autophagy of Lewy Body-Like α-Synuclein Inclusions. PLOS ONE 7, e52868 (2013).

41. S. Passaro, G. A.-O. Corso, J. A.-O. Wohlwend, M. A.-O. Reveiz, S. A.-O. Thaler, V. R. Somnath, N. Getz, T. A.-O. Portnoi, J. Roy, H. A.-O. X. Stark, D. Kwabi-Addo, D. A.-O. Beaini, T. A.-O. Jaakkola, R. Barzilay, Boltz-2: Towards Accurate and Efficient Binding Affinity Prediction. LID - 2025.06.14.659707 [pii] LID - 10.1101/2025.06.14.659707 [doi].

42. A. Ratti, V. Gumina, P. Lenzi, P. Bossolasco, F. Fulceri, C. Volpe, D. Bardelli, F. Pregnolato, A. Maraschi, F. Fornai, V. Silani, C. Colombrita, Chronic stress induces formation of stress granules and pathological TDP-43 aggregates in human ALS fibroblasts and iPSC-motoneurons. Neurobiology of Disease 145, 105051 (2020).

43. H. W. A. Cheng, T. B. Callis, A. A.-O. Montgomery, J. A.-O. Danon, W. T. Jorgensen, Y. A.-O. Ke, L. M. Ittner, E. L. Werry, M. Kassiou, Understanding In Vitro Pathways to Drug Discovery for TDP-43 Proteinopathies. LID - 10.3390/ijms232314769 [doi] LID - 14769.

44. H. Hartmann, W. Y. Ho, J. C. Chang, S.-C. Ling, Cholesterol dyshomeostasis in amyotrophic lateral sclerosis: cause, consequence, or epiphenomenon? The FEBS Journal, (2022).

45. S. M. Kim, M. Y. Noh, H. Kim, S. Y. Cheon, K. M. Lee, J. Lee, E. Cha, K. S. Park, K. W. Lee, J. J. Sung, S. H. Kim, 25-Hydroxycholesterol is involved in the pathogenesis of amyotrophic lateral sclerosis. Oncotarget 8, (2017).

46. J. Abdel-Khalik, E. Yutuc, P. J. Crick, J.-Å. Gustafsson, M. Warner, G. Roman, K. Talbot, E. Gray, W. J. Griffiths, M. R. Turner, Y. Wang, Defective cholesterol metabolism in amyotrophic lateral sclerosis. Journal of Lipid Research 58, 267–278 (2017).

47. J. C. Dodge, E. H. Jensen, J. Yu, S. P. Sardi, A. R. Bialas, T. V. Taksir, D. S. Bangari, L. S. Shihabuddin, Neutral Lipid Cacostasis Contributes to Disease Pathogenesis in Amyotrophic Lateral Sclerosis. Journal of Neuroscience, (202).

48. S. Michels, D. Kurz, A. Rosenbohm, R. S. Peter, S. Just, H. Bäzner, A. Börtlein, C. Dettmers, H.-J. Gold, A. Kohler, M. Naumann, P. Ratzka, A. C. Ludolph, D. Rothenbacher, G. Nagel, J. Dorst, A. L. S. R. S. S. G. the, Association of blood lipids with onset and prognosis of amyotrophic lateral sclerosis: results from the ALS Swabia registry. Journal of Neurology 270, 3082–3090 (2023).

49. D. Sapaly, F. Cheguillaume, L. Weill, Z. Clerc, O. Biondi, S. Bendris, C. Buon, R. Slika, E. Piller, V. K. Sundaram, A. da Silva Ramos, M. d. M. Amador, T. Lenglet, R. Debs, N. Le Forestier, P.-F. Pradat, F. Salachas, L. Lacomblez, A. Hesters, D. Borderie, D. Devos, C. Desnuelle, A.-S. Rolland, B. Periou, S. Vasseur, M. Chapart, I. Le Ber, A.-L. Fauret-Amsellem, S. Millecamps, T. Maisonobe, S. Leonard-Louis, A. Behin, F.-J. Authier, T. Evangelista, F. Charbonnier, G. Bruneteau, P. s. g. on behalf of the, Dysregulation of muscle cholesterol transport in amyotrophic lateral sclerosis. Brain 148, 788–802 (2025).

50. C. J. Kreple, S. Searles Nielsen, K. M. Schoch, T. Shen, M. Shabsovich, Y. Song, B. A. Racette, T. M. Miller, Protective Effects of Lovastatin in a Population-Based ALS Study and Mouse Model. Annals of Neurology 93, 881–892 (2023).

51. S. Luo, X. Wang, B. Ma, D. Liu, L. Li, L. Wang, N. Ding, L. Zou, J. Wang, J. Pan, D. Sang, H. Zhou, H. Qu, Y. Lu, L. Yang, Therapeutic potential of simvastatin in ALS: Enhanced axonal integrity and motor neuron survival through Apoa4 and Alb modulation.

52. J. R. Klim, L. A. Williams, F. Limone, I. Guerra San Juan, B. N. Davis-Dusenbery, D. A. Mordes, A. Burberry, M. J. Steinbaugh, K. K. Gamage, R. Kirchner, R. Moccia, S. H. Cassel, K. Chen, B. J. Wainger, C. J. Woolf, K. Eggan, ALS-implicated protein TDP-43 sustains levels of STMN2, a mediator of motor neuron growth and repair. Nature Neuroscience 22, 167–179 (2019).

53. M. Nolan, S. Aryal, I. S. Ndayambaje, M. Cao, P. Lee, M. Hovde, S. Yun, J. Wlaschin, A. Held, H. Beaussant, B. Wymann, C. Zong-Lee, S. M. Lim, X. Jiang, N. Ramesh, A. R. Agra Almeida Quadros, A. Boulos, N. Zinter, S. Salem, L. El-Tayar, M. Beccari, M. Presa, C. Jourdan Ferreras Reyes, Y. Y. Ruan, G. Griesman, C. Aguilar, J. Hawrot, H. Wheeler, Z. Melamed, B. P. Kleinstiver, M. Albers, D. W. Cleveland, R. E. Tanzi, C. M. Lutz, R. D. Hubbard, D. Kobayashi, M. Ward, C. R.R Alves, B. Wainger, C. L. Pichon, C. Lagier-Tourenne, Statins and genetic inhibition of the mevalonate pathway activate an ATF3-STMN2 regenerative program. bioRxiv, 2026.2002.2023.707492 (2026).

54. A. Rezaei, V. Kocsis-Jutka, Z. I. Gunes, Q. Zeng, G. Kislinger, F. Bauernschmitt, H. B. Isilgan, L. R. Parisi, T. Kaya, S. Franzenburg, J. Koppenbrink, J. Knogler, T. Arzberger, D. Farny, B. Nuscher, E. Katona, A. Dhingra, C. Yang, G. Gouna, K. D. LaClair, A. Janjic, W. Enard, Q. Zhou, N. Hagan, D. Ofengeim, E. Beltrán, O. Gokce, M. Simons, S. Liebscher, D. Edbauer, Correction of dysregulated lipid metabolism normalizes gene expression in oligodendrocytes and prolongs lifespan in female poly-GA C9orf72 mice. Nature Communications 16, 3442 (2025).

55. J. J. Yerbury, D. Gower, L. Vanags, K. Roberts, J. A. Lee, H. Ecroyd, The small heat shock proteins αB-crystallin and Hsp27 suppress SOD1 aggregation in vitro. Cell Stress and Chaperones 18, 251–257 (2013).

56. D. Cox, H. Ecroyd, The small heat shock proteins αB-crystallin (HSPB5) and Hsp27 (HSPB1) inhibit the intracellular aggregation of α-synuclein.

57. J. M. Gregory, T. P. Barros, S. Meehan, C. M. Dobson, L. M. Luheshi, The Aggregation and Neurotoxicity of TDP-43 and Its ALS-Associated 25 kDa Fragment Are Differentially Affected by Molecular Chaperones in Drosophila. PLOS ONE 7, e31899 (2012).

58. M. Feng, Y. Jin, S. Yang, A. M. Joachim, Y. Ning, L. M. Mori-Quiroz, J. Fromm, C. Perera, K. Zhang, K. A. Werbovetz, M. Z. Wang, Sterol profiling of Leishmania parasites using a new HPLC-tandem mass spectrometry-based method and antifungal azoles as chemical probes reveals a key intermediate sterol that supports a branched ergosterol biosynthetic pathway. International Journal for Parasitology: Drugs and Drug Resistance 20, 27–42 (2022).

59. V. Crippa, M. E. Cicardi, N. Ramesh, S. J. Seguin, M. Ganassi, I. Bigi, C. Diacci, E. Zelotti, M. Baratashvili, J. M. Gregory, C. M. Dobson, C. Cereda, U. B. Pandey, A. Poletti, S. Carra, The chaperone HSPB8 reduces the accumulation of truncated TDP-43 species in cells and protects against TDP-43-mediated toxicity.

60. L. T. Lin, A. Razzaq, S. E. Di Gregorio, S. Hong, B. Charles, M. H. Lopes, F. Beraldo, V. F. Prado, M. A. M. Prado, M. L. Duennwald, Hsp90 and its co-chaperone Sti1 control TDP-43 misfolding and toxicity.

61. R. Kuta, N. Larochelle, M. Fernandez, A. Pal, S. Minotti, M. Tibshirani, K. St Louis, B. J. Gentil, J. N. Nalbantoglu, A. Hermann, H. A.-O. Durham, Depending on the stress, histone deacetylase inhibitors act as heat shock protein co-inducers in motor neurons and potentiate arimoclomol, exerting neuroprotection through multiple mechanisms in ALS models. Cell Stress and Chaperones, (2020).

62. H. Shammas, C. Kloster Fog, P. Klein, A. Koustrup, M. T. Pedersen, A. S. Bie, T. Mickle, N. H. T. Petersen, T. Kirkegaard Jensen, S. Guenther, Mechanistic insights into arimoclomol mediated effects on lysosomal function in Niemann-pick type C disease. Molecular Genetics and Metabolism 145, 109103 (2025).

63. R. I. Morimoto, Regulation of the heat shock transcriptional response: cross talk between a family of heat shock factors, molecular chaperones, and negative regulators. Genes & development 12, 3788–3796 (1998).

64. D. R. Sojka, A. Gogler-Pigłowska, N. Vydra, A. J. Cortez, P. T. Filipczak, Z. Krawczyk, D. A.-O. Scieglinska, Functional redundancy of HSPA1, HSPA2 and other HSPA proteins in non-small cell lung carcinoma (NSCLC); an implication for NSCLC treatment.

65. J. Chua, H. De Calbiac, E. Kabashi, S. Barmada, Autophagy and ALS: mechanistic insights and therapeutic implications. Autophagy, (2022).

66. L. A.-O. Hayes, P. Kalab, Emerging Therapies and Novel Targets for TDP-43 Proteinopathy in ALS/FTD.

67. L. Chen, E. H. J. Krekels, P. E. Verweij, J. B. Buil, C. A. J. Knibbe, R. J. M. Brüggemann, Pharmacokinetics and Pharmacodynamics of Posaconazole. Drugs 80, 671–695 (2020).

68. A. K. Gupta, D. C. Lyons, The Rise and Fall of Oral Ketoconazole.

69. P. Pitisuttithum, J. R. Negroni R Fau-Graybill, B. Graybill Jr Fau-Bustamante, P. Bustamante B Fau-Pappas, S. Pappas P Fau-Chapman, R. S. Chapman S Fau-Hare, C. J. Hare Rs Fau-Hardalo, C. J. Hardalo, Activity of posaconazole in the treatment of central nervous system fungal infections. Journal of Antimicrobial Chemotherapy, (2005).

70. J. D. Thompson, T. J. Gibson, D. G. Higgins, Multiple sequence alignment using ClustalW and ClustalX. Current Protocols in Bioinformatics, (2002).

71. K. J. Livak, T. D. Schmittgen, Analysis of relative gene expression data using real-time quantitative PCR and the 2(-Delta Delta C(T)) Method. Methods 25, (2001).

72. R. A. Friesner, J. L. Banks, R. B. Murphy, T. A. Halgren, J. J. Klicic, D. T. Mainz, M. P. Repasky, E. H. Knoll, M. Shelley, J. K. Perry, D. E. Shaw, P. Francis, P. S. Shenkin, Glide: a new approach for rapid, accurate docking and scoring. 1. Method and assessment of docking accuracy. Journal of medicinal chemistry 47, 1739–1749 (2004).

73. M. J. Workman, R. G. Lim, J. Wu, A. Frank, L. Ornelas, L. Panther, E. Galvez, D. Perez, I. Meepe, S. Lei, V. Valencia, E. Gomez, C. Liu, R. Moran, L. Pinedo, S. Tsitkov, R. Ho, J. A. Kaye, T. Thompson, J. D. Rothstein, S. Finkbeiner, E. Fraenkel, D. Sareen, L. M. Thompson, C. N. Svendsen, Large-scale differentiation of iPSC-derived motor neurons from ALS and control subjects. Neuron, (2023).

74. P. Walsh, V. Truong, S. Nayak, M. Saldías Montivero, W. C. Low, A. M. Parr, J. R. Dutton, Accelerated differentiation of human pluripotent stem cells into neural lineages via an early intermediate ectoderm population. Stem Cells, (2020).

75. C. B. Pantazis, A. Yang, E. Lara, J. A. McDonough, C. Blauwendraat, L. Peng, H. Oguro, J. Kanaujiya, J. Zou, D. Sebesta, G. Pratt, E. Cross, J. Blockwick, P. Buxton, L. Kinner-Bibeau, C. Medura, C. Tompkins, S. Hughes, M. Santiana, F. Faghri, M. A. Nalls, D. Vitale, S. Ballard, Y. A. Qi, D. M. Ramos, K. M. Anderson, J. Stadler, P. Narayan, J. Papademetriou, L. Reilly, M. P. Nelson, S. Aggarwal, L. U. Rosen, P. Kirwan, V. Pisupati, S. L. Coon, S. W. Scholz, T. Priebe, M. Öttl, J. Dong, M. Meijer, L. J. M. Janssen, V. S. Lourenco, R. van der Kant, D. Crusius, D. Paquet, A. C. Raulin, G. Bu, A. Held, B. J. Wainger, R. M. C. Gabriele, J. M. Casey, S. Wray, D. Abu-Bonsrah, C. L. Parish, M. S. Beccari, D. W. Cleveland, E. Li, I. V. L. Rose, M. Kampmann, C. Calatayud Aristoy, P. Verstreken, L. Heinrich, M. Y. Chen, B. Schüle, D. Dou, E. L. F. Holzbaur, M. C. Zanellati, R. Basundra, M. Deshmukh, S. Cohen, R. Khanna, M. Raman, Z. S. Nevin, M. Matia, J. Van Lent, V. Timmerman, B. R. Conklin, K. Johnson Chase, K. Zhang, S. Funes, D. A. Bosco, L. Erlebach, M. Welzer, D. Kronenberg-Versteeg, G. Lyu, E. Arenas, E. Coccia, L. Sarrafha, T. Ahfeldt, J. C. Marioni, W. C. Skarnes, M. R. Cookson, M. E. Ward, F. T. Merkle, A reference human induced pluripotent stem cell line for large-scale collaborative studies. Cell Stem Cell, (2022).

