## Supplemental Information for "From anti-fungal to potential neurotherapeutic: Posaconazole as an effective inhibitor of cellular TDP-43 pathology"

**Title**

This file includes:

- Figs. S1-S13

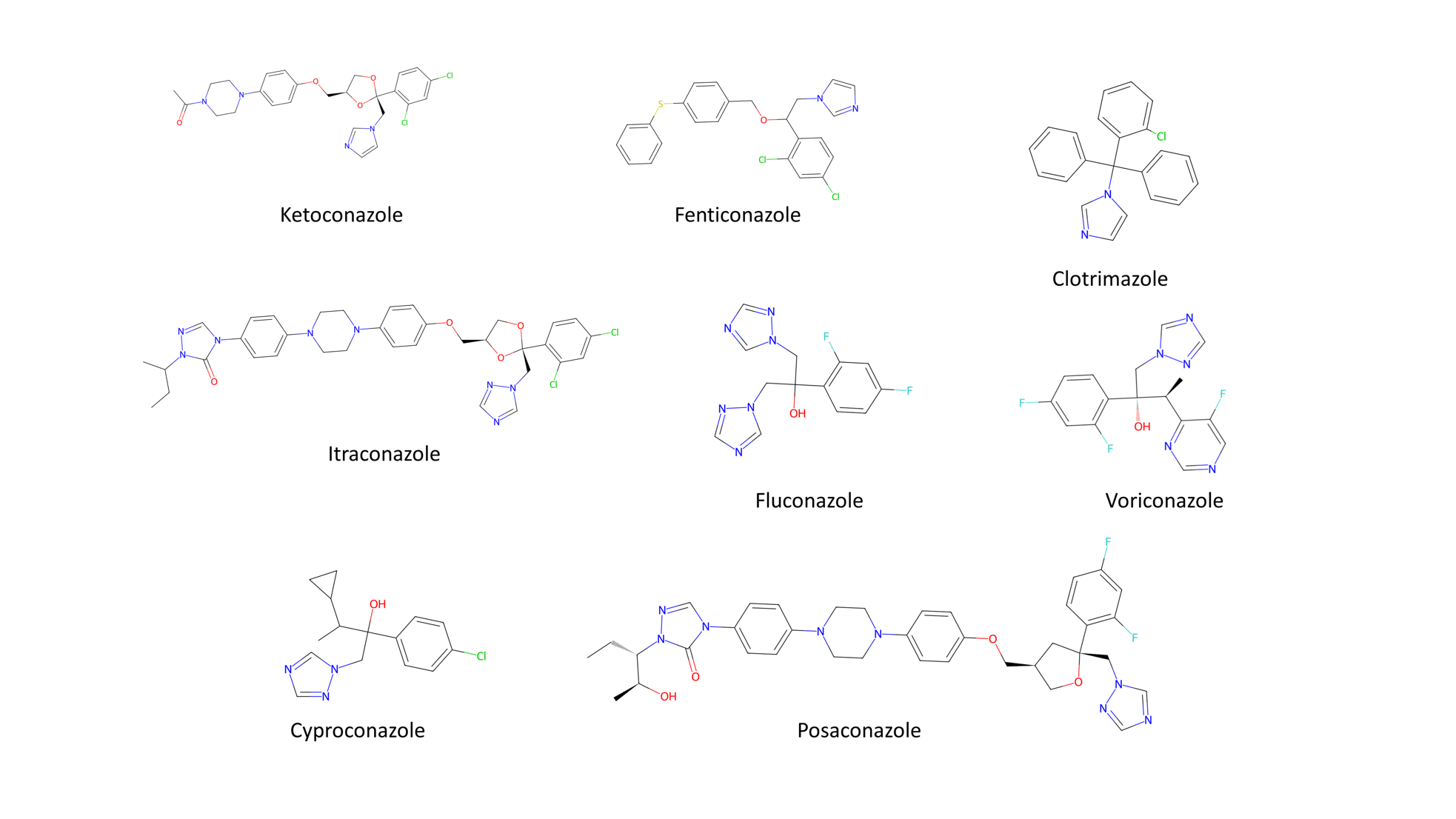

**Fig. S1. Chemical structures of the 8 azole-based CYP51 inhibitors tested in this study.**

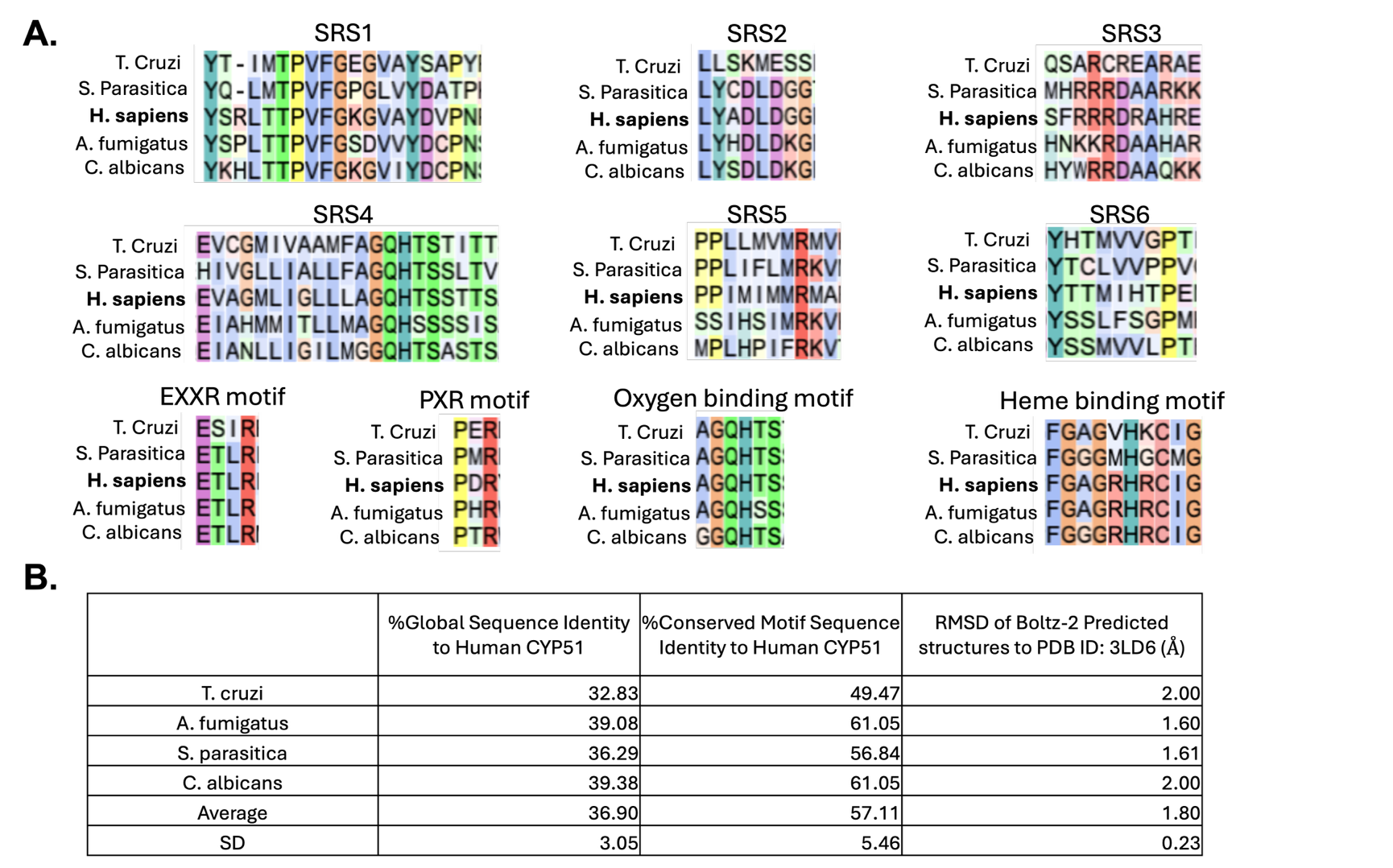

**Fig. S2. Multiple sequence alignment at conserved motifs of fungal CYP51 with experimental azole data and human CYP51. (A)** Multiple sequence alignment of conserved motifs between human and fungal CYP51 with azole experimental data shown in Fig. 1A. Motifs included in the analysis are the substrate recognition sites 1-6 (SRS1-6) involved in lanosterol binding, EXXR and PXR motifs which form an E-R-R triad responsible for stabilizing the structure near the heme binding pocket, oxygen binding motif responsible for activating oxygen for the catalytic function of the enzyme and the heme binding motif responsible for coordinating the prosthetic heme group crucial for catalytic activity (*1*). **(B)** Summary of global and conserved motif sequence identity (defined as identical residues) between fungal CYP51s and human CYP51, as well as RMSDs from structural alignments of Boltz-2 predicted structures of fungal CYP51s to human CYP51 (PDB ID: 3LD6).

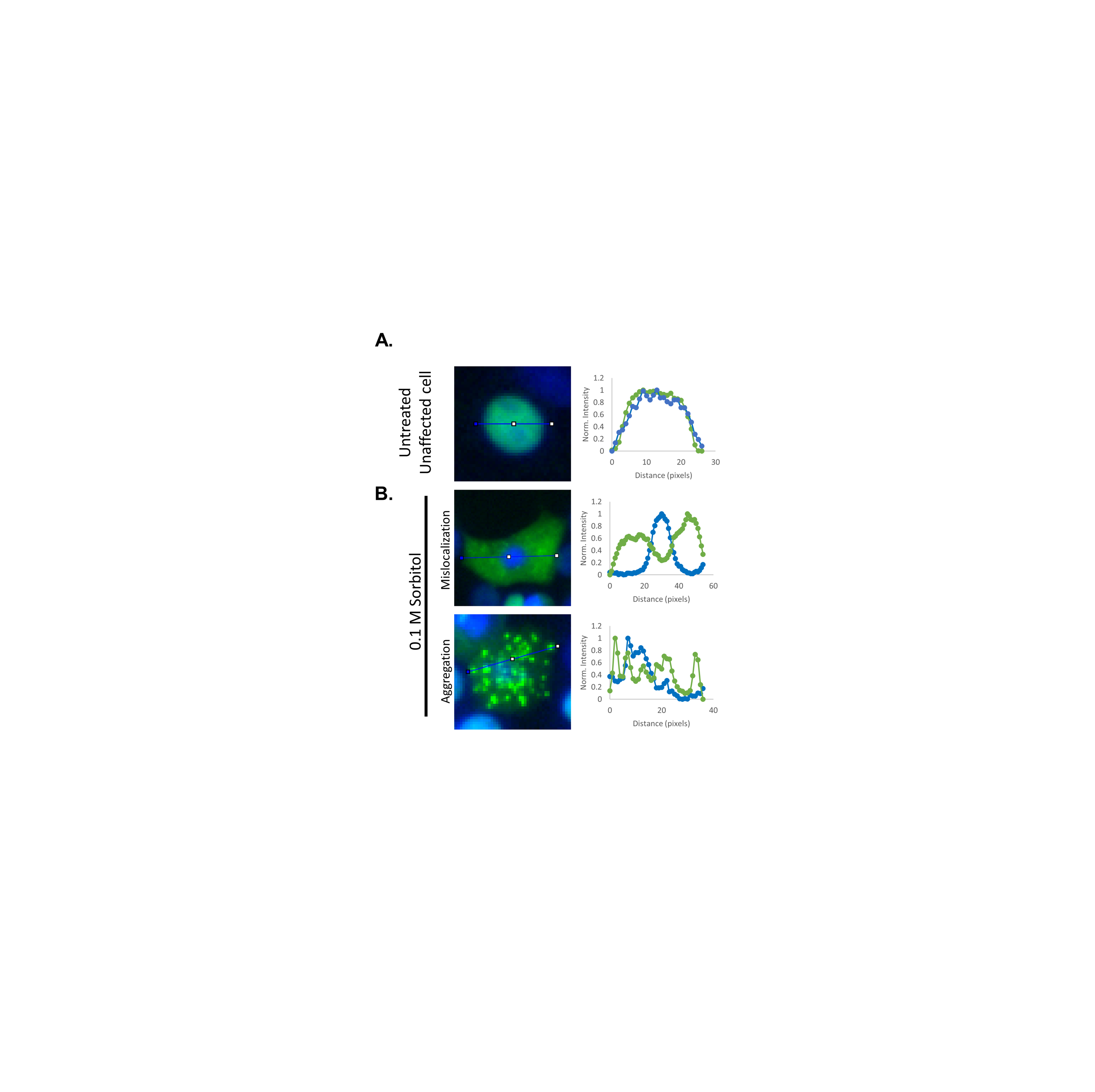

**Fig. S3. Summary of sorbitol-induced TDP-43 mislocalization and aggregation model in HEK293T cells. (A)** Representative TDP-43-mNg and nuclear stain image of untreated HEK293T cell with accompanying line intensity plot. **(B)** Representative TDP-43-mNg and nuclear stain image of 0.1 M sorbitol treated HEK293T cells with accompanying line intensity plots. Sorbitol induces both cytoplasmic mislocalization and puncta formation.

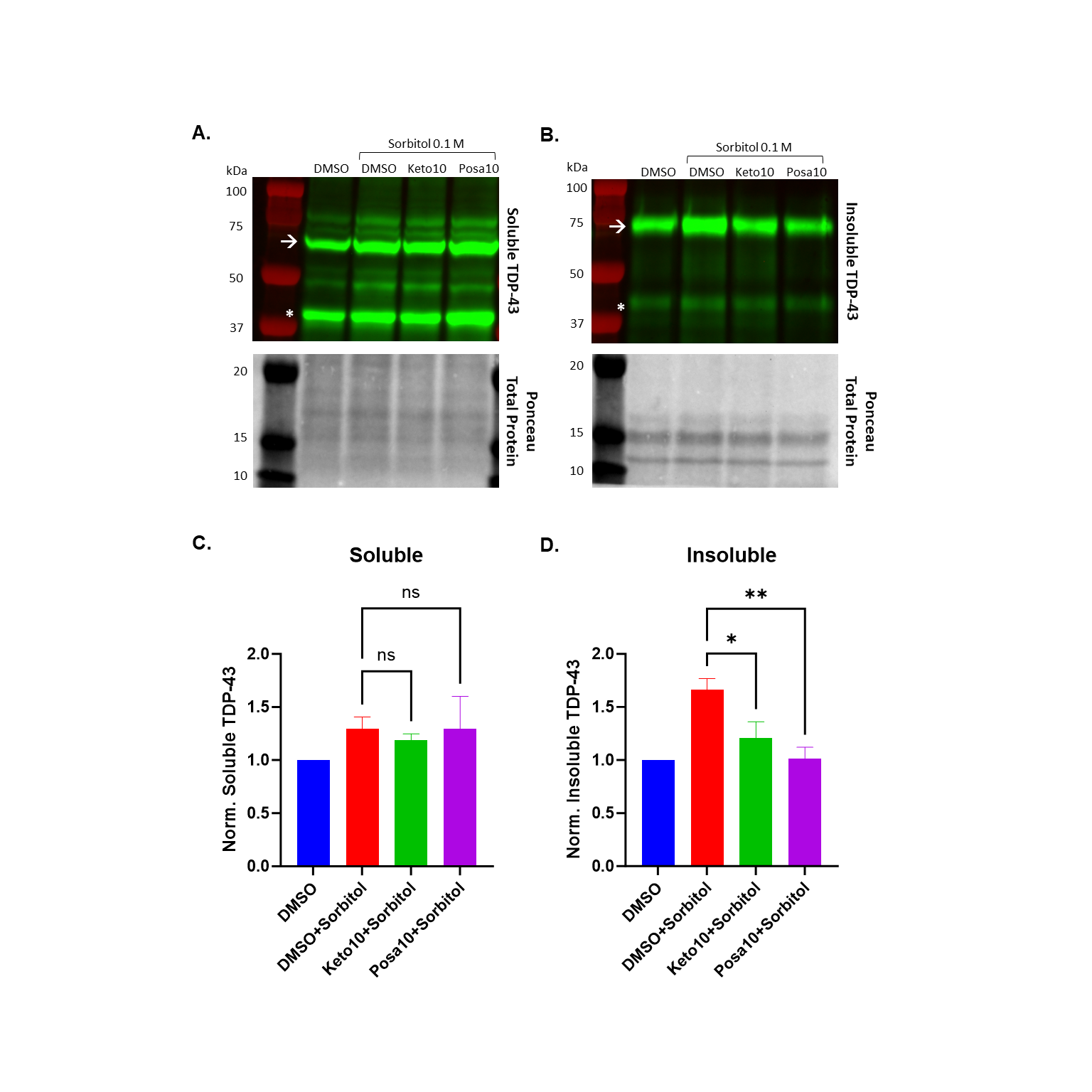

**Fig. S4. Soluble/insoluble protein fraction quantification.** Quantification of **(A)** soluble and **(B)** insoluble TDP-43 protein (insoluble fraction data repeated for ease of comparison). Data shown are mean ± SEM of N=5 independent experiments, analyzed via a one-way ANOVA with Bonferroni correction for multiple comparisons relative to DMSO/sorbitol-treated cells (red bar, *p < 0.05, **p < 0.01).

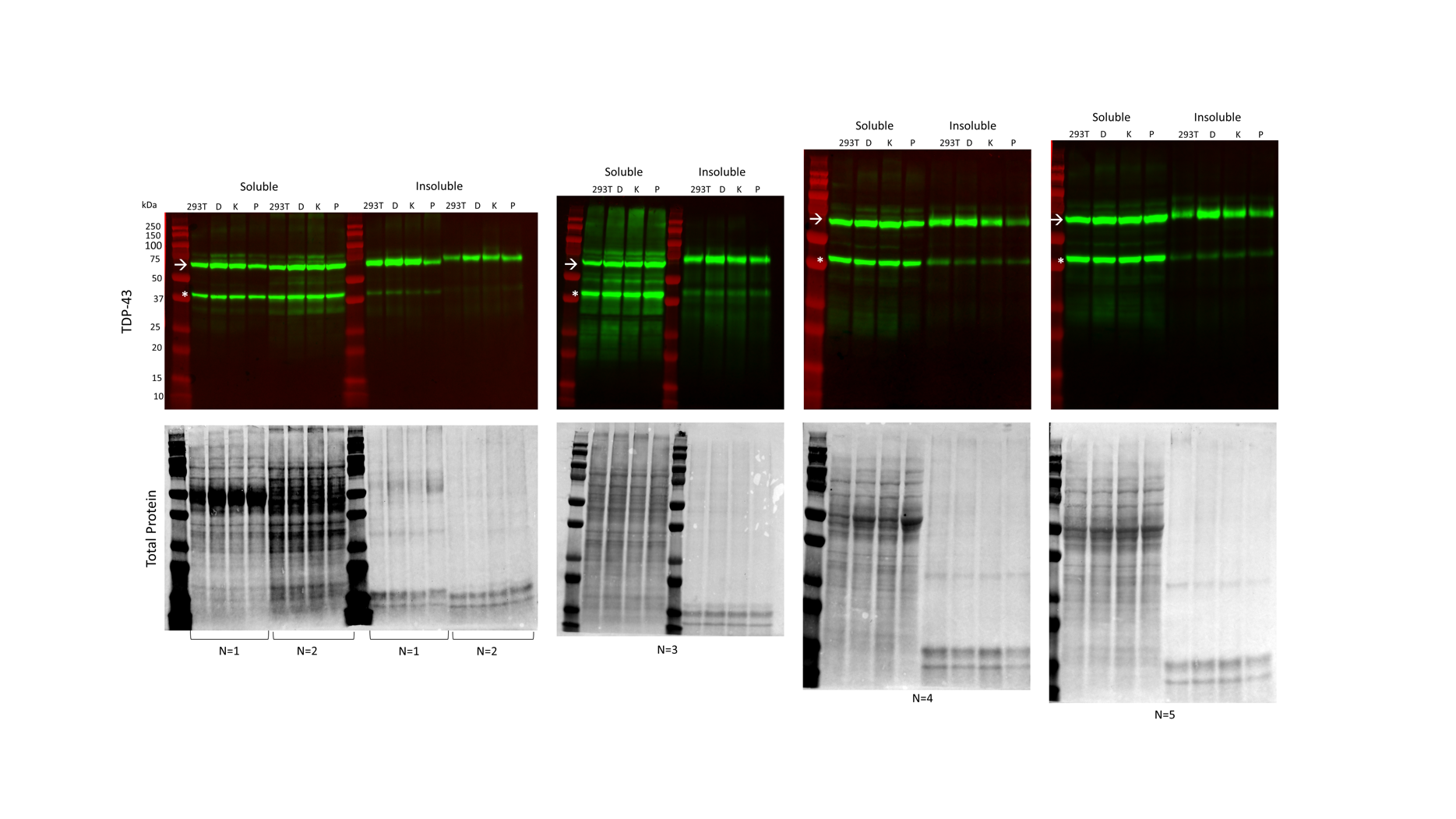

**Fig. S5. Soluble/insoluble protein fraction western blots.** White asterisk (*) indicates endogenous TDP-43, white arrow (🡪) indicates TDP-43-mNg (which was used for quantification). Note that replicates N=1 and N=2 were run in the same gel and grouped by soluble and insoluble fractions.

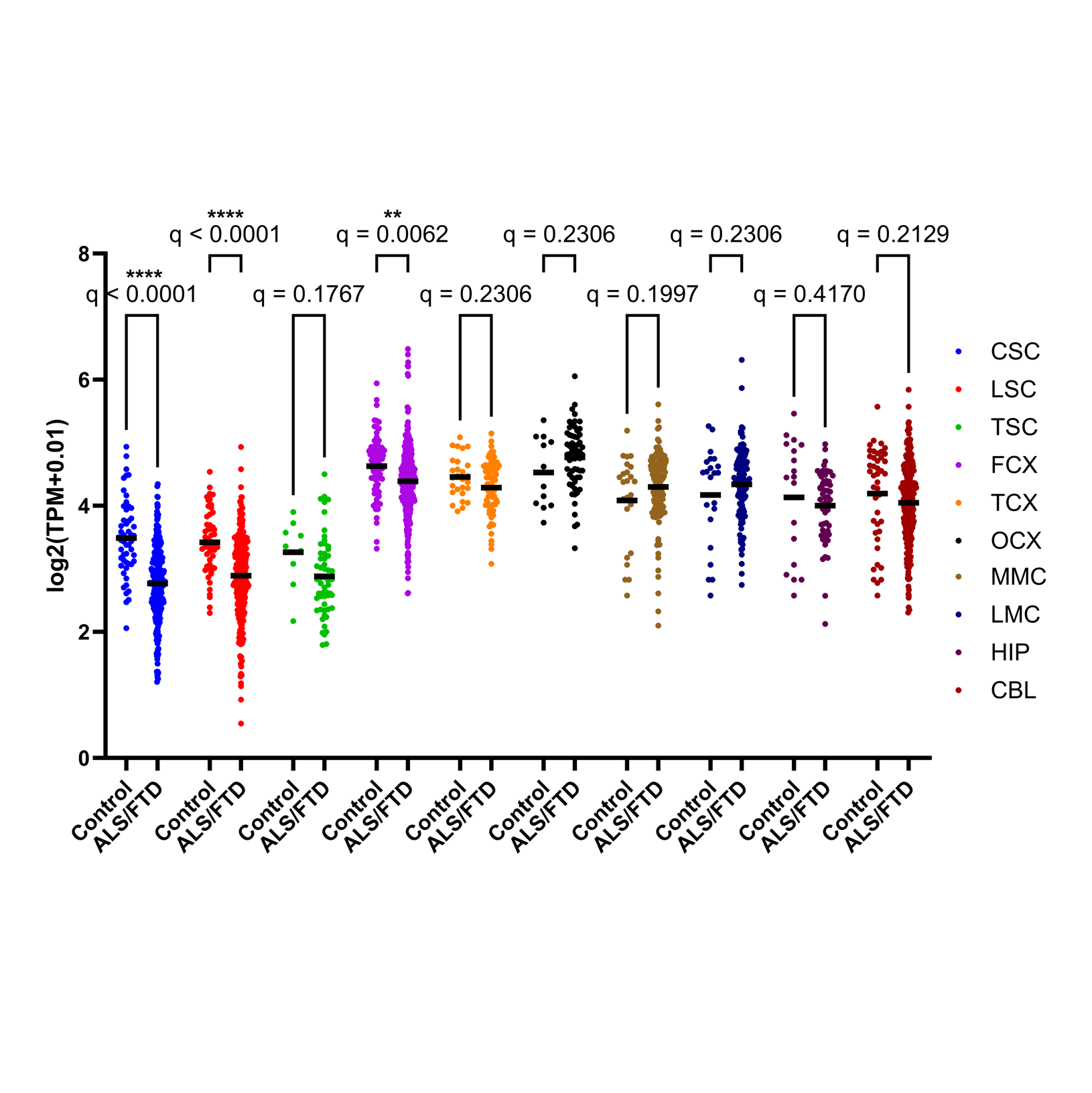

**Fig. S6. Expression of SREBP2 across several tissue types characterized in the NYGC ALS Consortium dataset. (A)** Transcripts per million (TPM) counts for SREBP2 mRNA in tissues of healthy controls and ALS/FTD patients extracted from the NYGC ALS Consortium dataset (*2*). Data was analyzed via a two-way ANOVA with Benjamini-Hochberg False Discovery Rate (FDR) correction (**q<0.01, ****q < 0.0001, q-values equate to FDR-adjusted p-values).

CSC = Cervical Spinal Cord

LSC = Lumbar Spinal Cord

TSC = Thoracic Spinal Cord

FCX = Frontal Cortex

TCX = Temporal Cortex

OCX = Occipital Cortex

MMC = Medial Motor Cortex

LMC = Lateral Motor Cortex

HIP = Hippocampus

CBL = Cerebellum

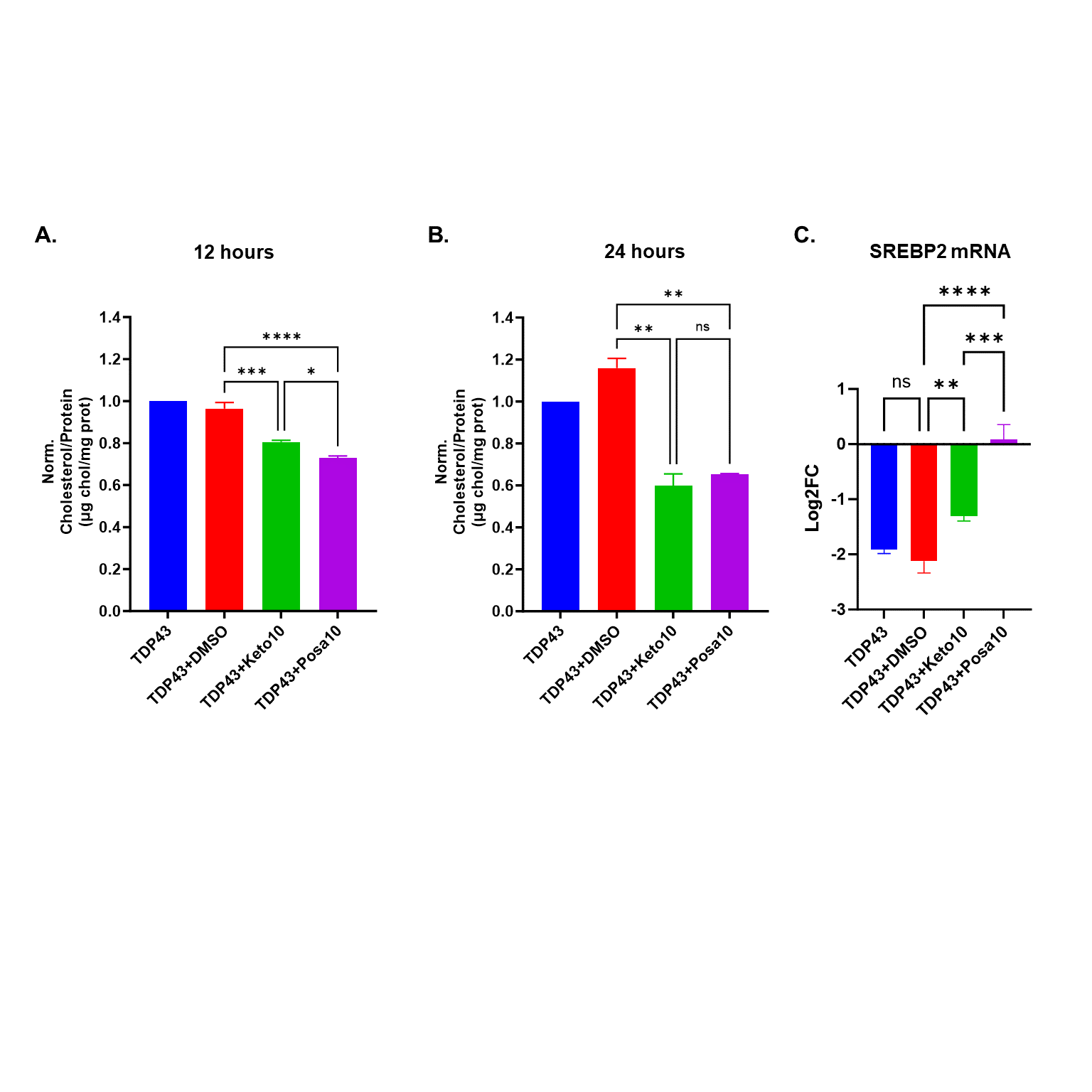

**Fig. S7. Cellular cholesterol levels under TDP-43 overexpression with 10 μM** **drug treatment and 100 μM** **cholesterol supplementation at 12 and 24 hours of treatment.** Data shown are mean ± SEM of N=3 independent experiments, analyzed via a one-way ANOVA with Bonferroni correction for multiple comparisons (**p < 0.01, ***p < 0.001).

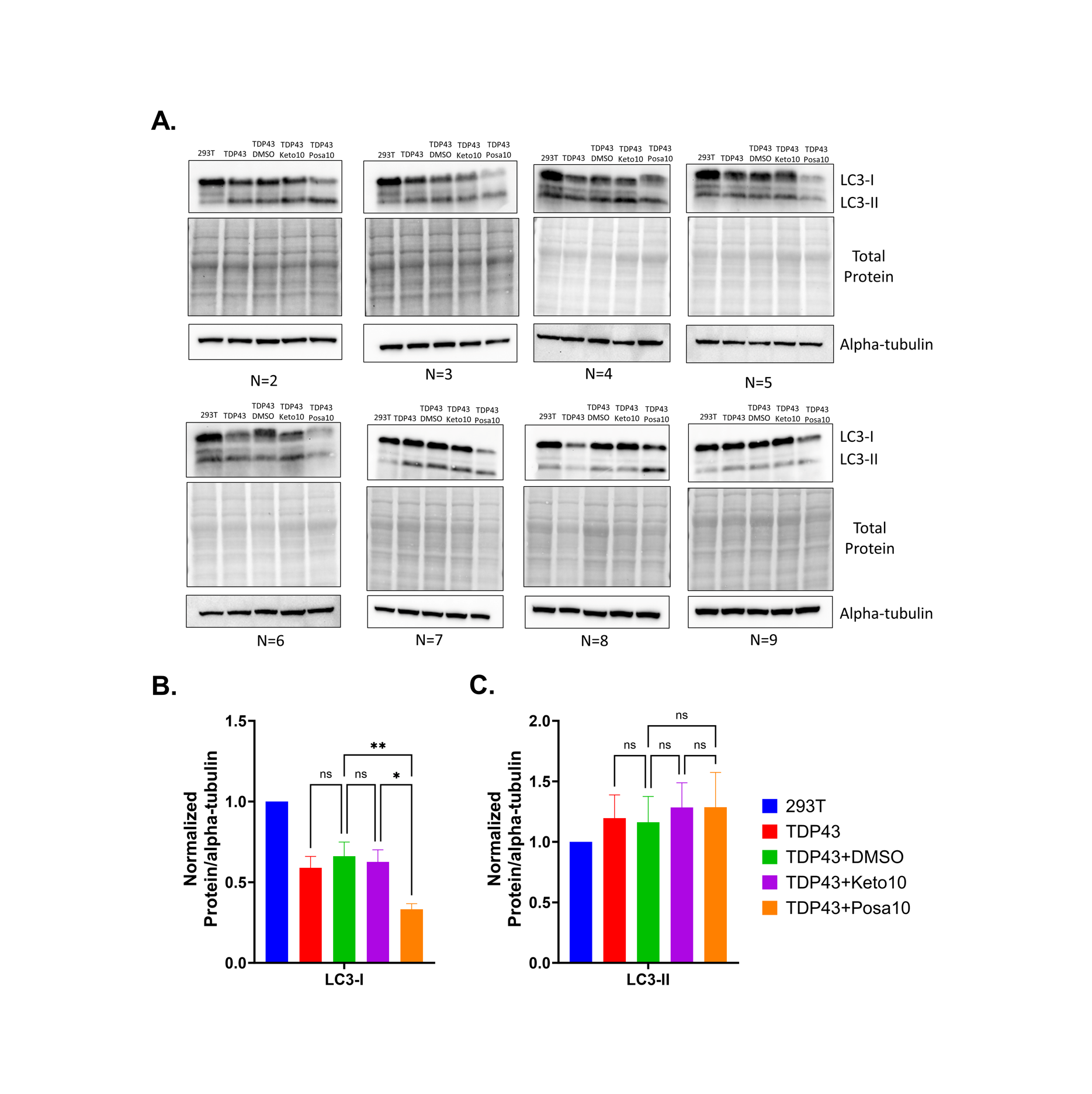

**Fig. S8. Western blots (N=2-9) probing LC3-I and LC3-II in TDP-43 overexpressing cells treated with 10 μM** **azoles.** **(A)** Western blots of LC3, Ponceau S (total protein) alpha-tubulin loading controls. **(B-C)** Western blot quantification of LC3-I and LC3-II. Data shown are mean ± SEM of N=9 independent experiments, analyzed via a one-way ANOVA with Bonferroni correction for multiple comparisons relative to DMSO treated cells (*p < 0.05, **p < 0.01).

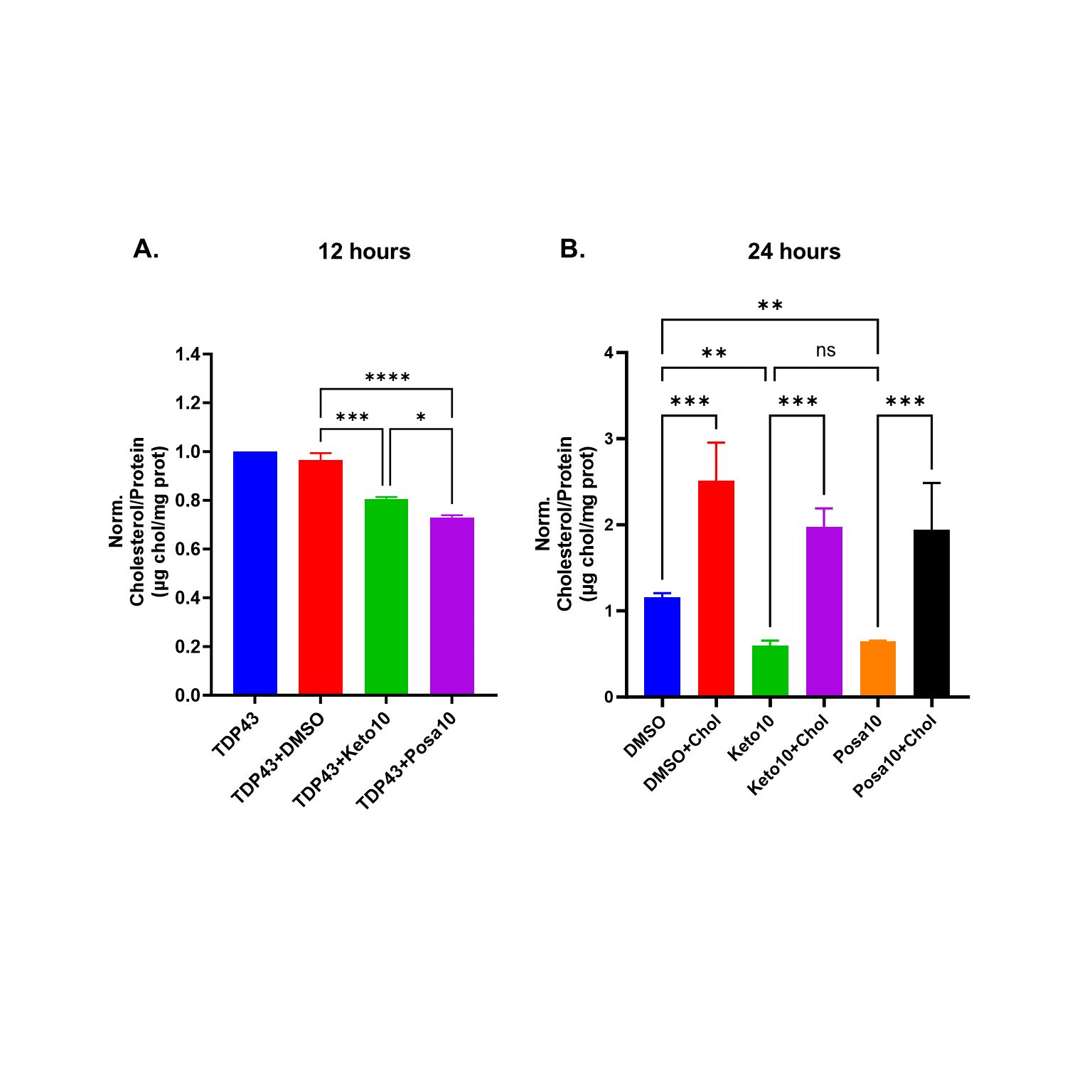

**Fig. S9. Cholesterol supplementation validation in HEK293T cells.** Cellular cholesterol levels normalized to total protein for DMSO, 10 μM ketoconazole and 10 μM posaconazole -/+ 100 μM supplemented cholesterol treated for 24 hours. Data shown are mean ± SEM of N=3 independent experiments, analyzed via a one-way ANOVA with Bonferroni correction for multiple comparisons (**p < 0.01, ***p < 0.001).

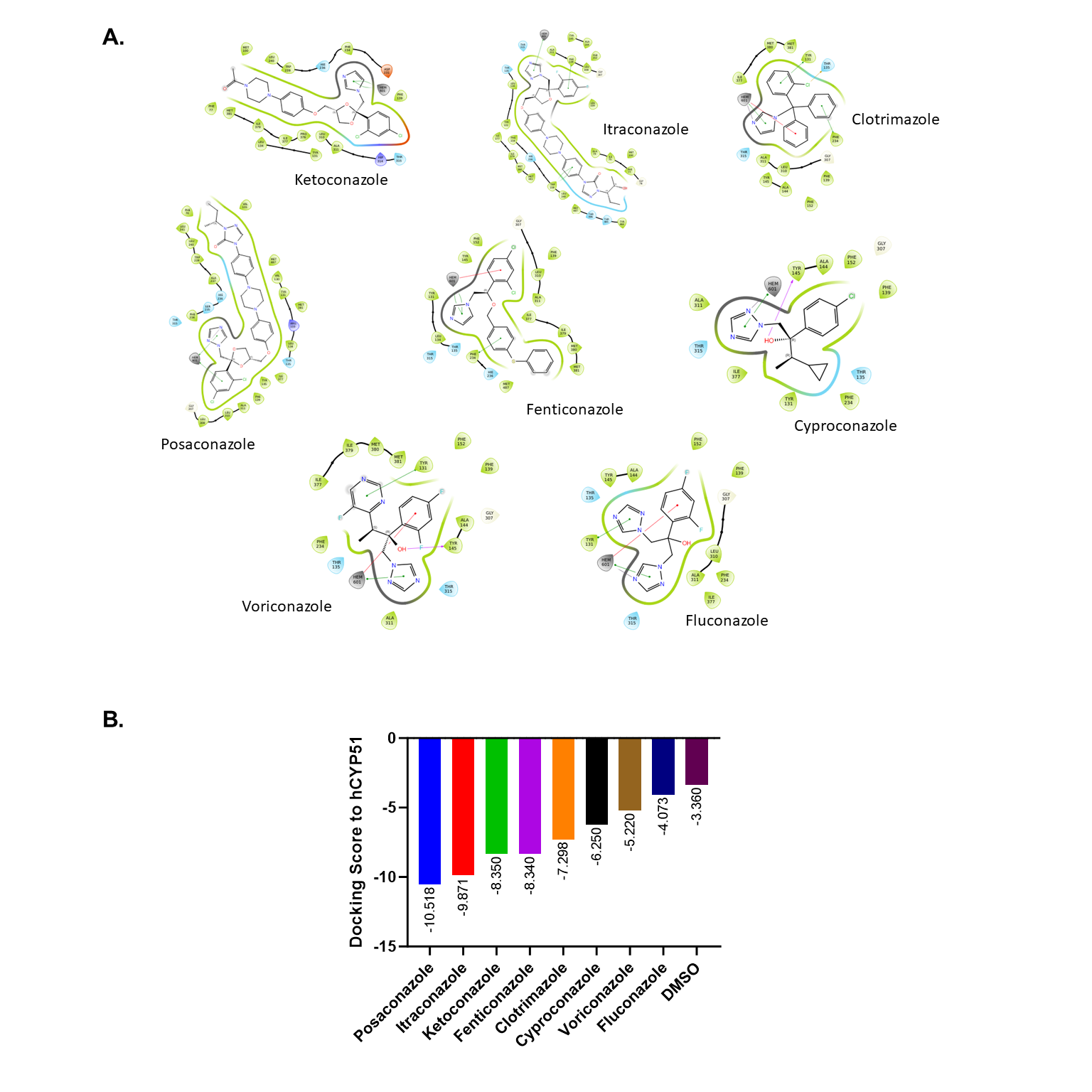

**Fig. S10. Protein-ligand interaction diagrams of inhibitors with human CYP51.** Maestro-generated protein ligand interactions for metal coordination-constrained docking campaign against human CYP51 (PDB ID: 3LD6). Note that all docked poses display the known azole-heme binding mode interactions for this class of inhibitors.

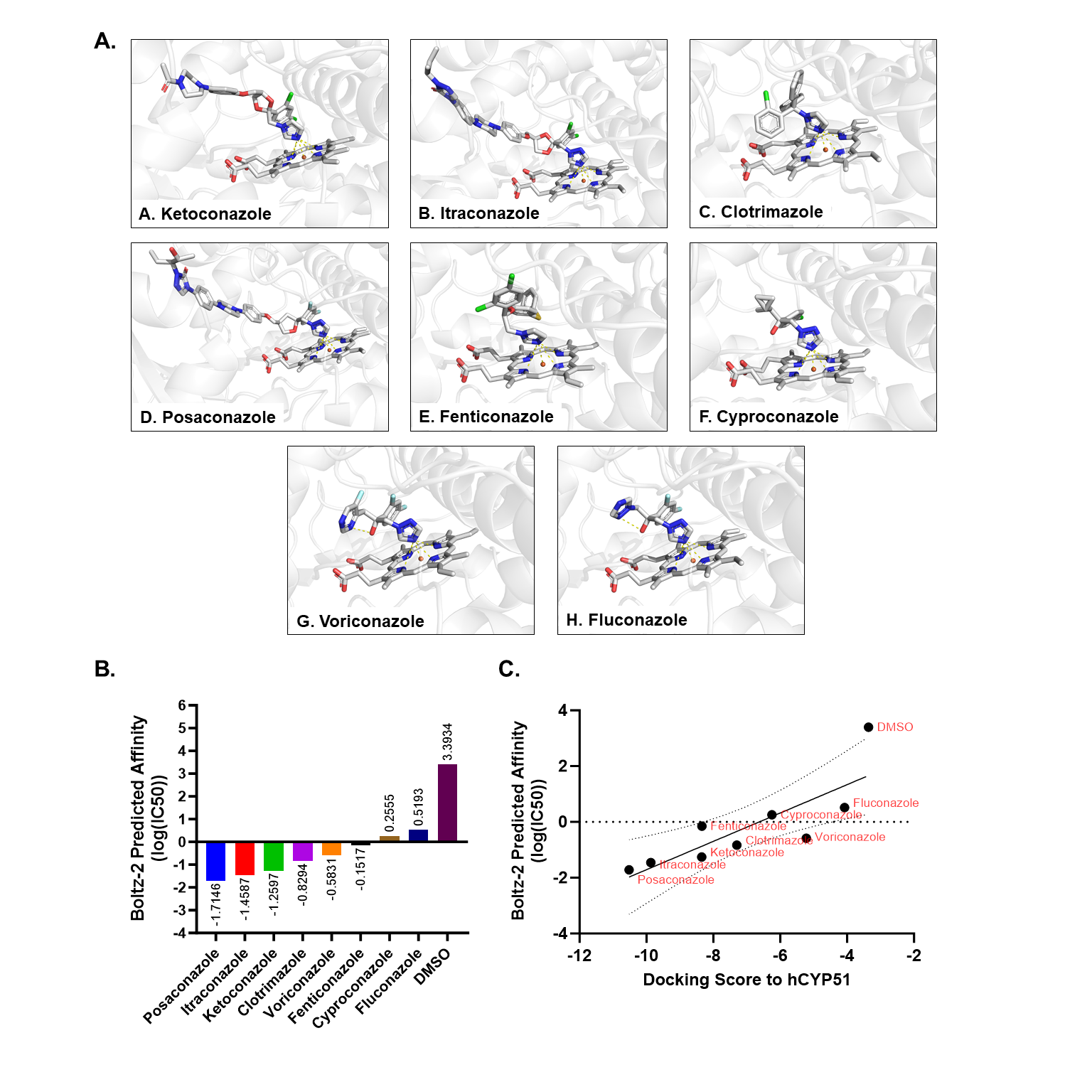

**Fig. S11. Validation of molecular docking against human CYP51 using Boltz-2. (A)** Boltz-2 structure predictions for azoles and human CYP51. Yellow dashed lines indicate polar contacts between inhibitors and heme. **(C)** Correlation between Boltz-2 predicted affinities (log(IC50)) and Glide docking scores (p = 0.0069).

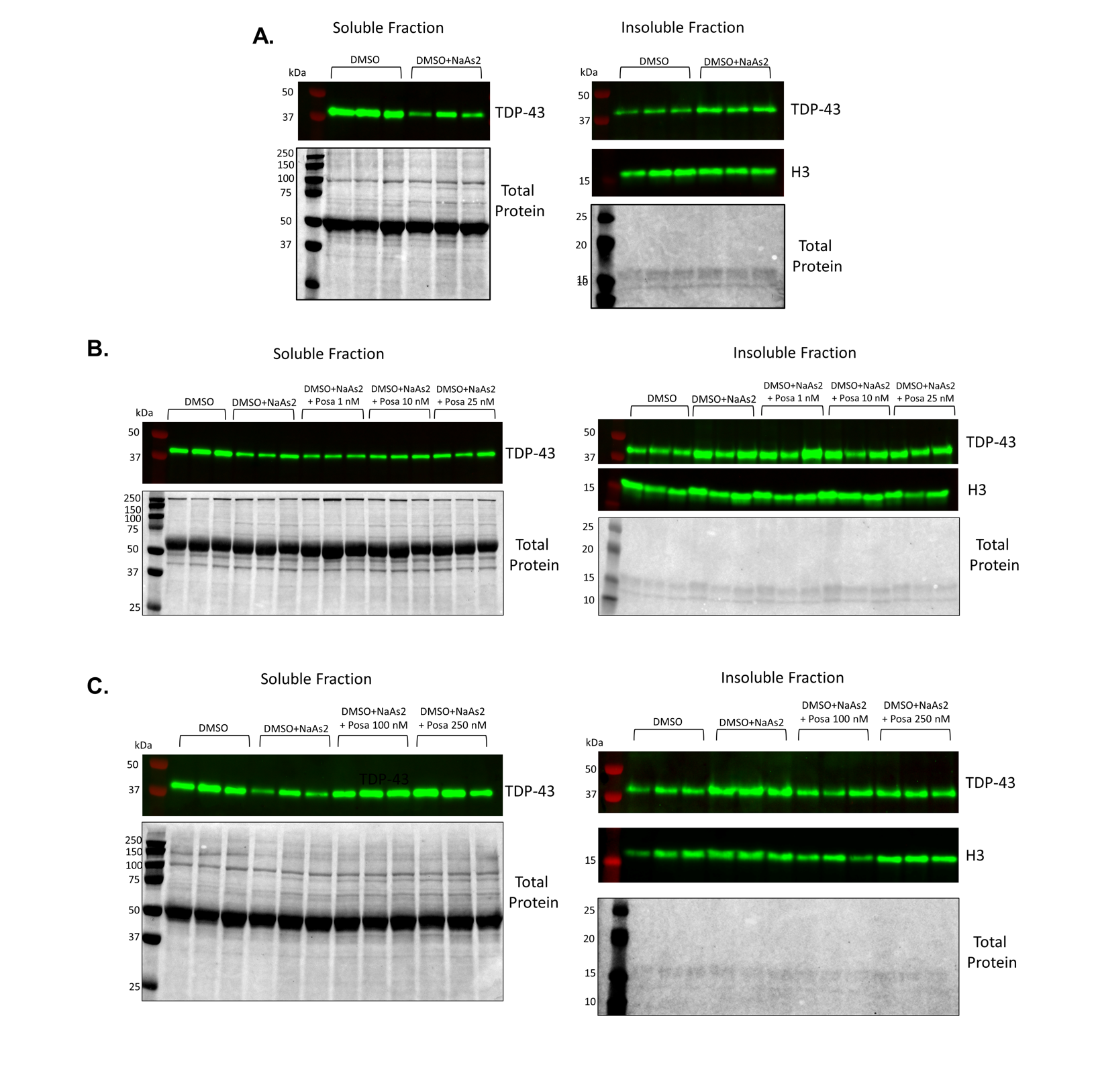

**Fig. S12. Western blots of low dose NaAs2 motor neuron model with DMSO and posaconazole treatment. (A)** N=1-3 soluble and insoluble fraction blots for DMSO -/+ NaAs2 treatments. **(B)** N=4-6 (for DMSO -/+ NaAs2) and N=1-3 (for NaAs2 + 1, 10 and 25 nM posaconazole) soluble and insoluble blots. **(C)** N=7-9 (for DMSO -/+ NaAs2) and N=1-3 (for NaAs2 + 100 and 250 nM posaconazole) soluble and insoluble blots.TDP-43 and total protein (Ponceau S, used for normalization) were probed for soluble fraction. TDP-43, total protein (Ponceau S) and histone 3 (H3, used for normalization) were probed for insoluble fraction. Data shown are mean ± SEM of N=3-9 independent treatments, analyzed via a one-way ANOVA with Bonferroni correction for multiple comparisons relative to DMSO/NaAs2 treated cells (*p < 0.05, **p < 0.01, ***p < 0.001).

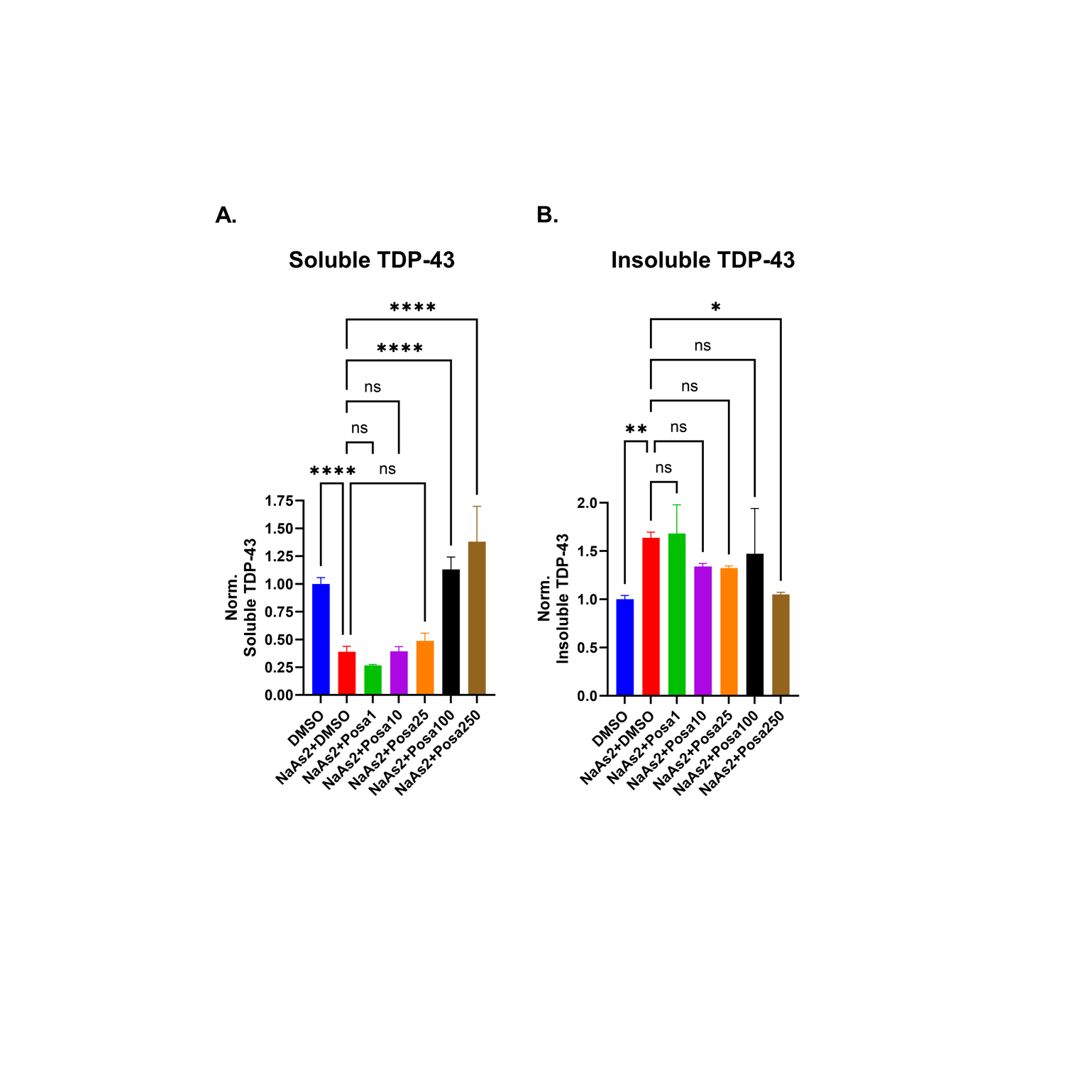

**Fig. S13. Western blot quantification of soluble and insoluble TDP-43 levels in motor neurons. (A)** Normalized soluble TDP-43 levels extracted from motor neurons treated with NaAs2 and posaconazole titration. **(B)** Normalized insoluble TDP-43 levels extracted from motor neurons treated with NaAs2 and posaconazole titration. Ratio of quantifications in (A) and (B) is the insoluble/soluble ratio shown in main Fig. 6C. Data shown are mean ± SEM of N=3-9 independent treatments, analyzed via a one-way ANOVA with Bonferroni correction for multiple comparisons relative to DMSO/NaAs2 treated cells (*p < 0.05, **p < 0.01, ****p < 0.0001).

**References**

1.	B. Celia-Sanchez, B. Mangum, M. Brewer, M. Momany, Analysis of Cyp51 protein sequences shows 4 major Cyp51 gene family groups across fungi. *G3: Genes, Genomes, Genetics*, (2022).

2.	J. Humphrey, A. Oku, M. Byrska-Bishop, A. O. Basile, U. S. Evani, A. Corvelo, A. Tokolyi, K. Bp, A. Réal, Y. Kim, M. L. Bond, W. E. Clarke, R. Fu, H. Geiger, S. Chang, T. Naito, B. Jang, R. Musunuri, W. H. Dredge, R. Al-Abri, B. N. Hoover, D. Manaa, J. McClintock, F. P. Singh, M. H. Pedersen, A. Runnels, N. Propp, S. Fennessey, H.-H. Won, M. C. Zody, G. Narzisi, N. Robine, T. Lappalainen, D. Fagegaltier, G. Gürsoy, D. A. Knowles, T. Raj, N. A. Consortium, M. B. Harms, H. Phatnani, The New York Genome Center ALS Consortium resource integrates postmortem tissue transcriptomics and whole genome sequencing to empower biological discovery. *medRxiv*, 2026.2004.2029.26350889 (2026).
